# Multi-stage efficient coding of perception and value in goal-directed behavior

**DOI:** 10.64898/2026.06.08.730978

**Authors:** Saurabh Bedi, Gilles de Hollander, Maximilian Harl, Christian C. Ruff

## Abstract

To act effectively, the brain must transform information through a chain of processing stages, from sensing the environment, to evaluating options, to selecting actions. Because neural resources are limited, each stage should represent information efficiently. Yet efficient coding has been studied almost exclusively in perception, and always one stage at a time. Whether perception and valuation are each governed by their own efficient code, and how these codes interact along the pathway from sensation to action, remains an open question. We developed a formal framework, comparing models in which efficient coding and Bayesian decoding shape perception only, valuation only, or both. To tease apart contributions from each stage, we designed an experiment that independently varied how stimuli map onto values. In a preregistered study, behavior was best explained by efficient coding operating at both stages, with each stage tracking its own objective prior. Crucially, changing the value distribution reversed the classic repulsion biases seen in orientation perception, revealing separable efficient codes in perception and valuation. The two stages also operated on different timescales: perceptual representations reflected stable, long-term environmental structure, while value representations updated rapidly with changing context. Together, these findings show that separable efficient codes at successive processing stages combine to shape behavior. This principle likely extends well beyond perception and valuation: whenever behavior depends on abstract, constructed representations rather than raw sensory signals, the brain may solve the efficiency problem anew at each stage.

## 1 Introduction

Adaptive behavior requires the brain to transform sensory input into flexible, goal-dependent representations that guide action [1]. Identical sensory inputs can drive very different behavior depending on current goals. For example, green bananas are attractive for storing but not for eating, whereas yellow bananas are attractive for eating but not for storing. This goal-dependent behavior is thought to arise from abstract value or utility variables[2–5] that the brain actively constructs from perceptual inputs depending on current goals [1, 2, 6–9]. Such abstract representations have been proposed to underlie higher-order cognition across many domains, from planning and concept learning to social cognition and cognitive control [10–15]. Neural evidence suggests that they are organized hierarchically and become progressively more abstract and task-dependent across successive processing stages [16–18]. Crucially, each of these representations must be maintained by a noisy [19] and resource-limited brain [20– 22]. This raises a central question: when information is transformed across multiple representational stages, how do noise and resource limitations jointly shape those representations and the systematic behavioral biases, distortions, and variability they produce? Here we address this question in the tractable case of value-based decision-making, where both the perceptual input and the constructed value representation can be expressed as scalar quantities.

Systematic behavioral biases offer a window into the efficient codes that underlie decision-making: by asking where biases originate and what statistics they reflect, we can infer how the brain represents information at each stage, from perception to action. Within perceptual and value-based decision-making, different traditions have exploited this logic, locating biases at different stages from perception to action and attributing them to different mechanisms. In perception, biases are often modeled as arising from Bayesian decoding of noisy sensory representations under prior expectations [23–26], and from efficient coding of those representations under limited resources, with the neural code adapting to environmental statistics, leading to predictable biases [27–30]. For value-based decisions, value representations are themselves sensitive to contextual statistics, exhibiting range adaptation and divisive normalization with respect to experienced value distributions [31–34] — signatures consistent with efficient coding operating in value space [35–37]. Recent work has begun to bridge these traditions by showing that representational stages are not independent but may influence each other [38]: perceptual distortions from noise and resource limits may propagate into downstream economic decisions [39–42], while task goals and values may in turn reshape upstream sensory representations [43]. Yet it remains unresolved what mechanisms govern each stage separately, whether the same computational principles operate across stages simultaneously, and how distortions within stages and in the mappings between them jointly shape behavior. Existing accounts still largely attribute distortions to a single locus, leaving the combined influence of noise and efficient coding across the full multi-stage process largely unexamined. This leaves open a recently emphasized question: at which levels of the neural processing stream, and over which temporal scales, should efficient adaptations shape information for behavior [44]?

A central challenge in determining where distortions arise along the perception–valuation pathway is that values are not intrinsic properties of the world but are constructed from perceptual inputs through context- and goal-dependent mappings. Notably, because subjective value representations are derived from perceptual features under a given goal, the statistical structure of values inherits the statistical structure of perceptual inputs. For example, if value increases with banana ripeness, an environment containing mostly ripe yellow bananas necessarily also contains many high-value items, whereas an environment dominated by unripe green bananas contains mostly low-value items. Perceptual and value distributions are therefore intrinsically coupled, so that changing perceptual input statistics inevitably reshapes value statistics for a fixed mapping/goal (Fig. 1b). This creates an identification problem. While prior work suggests that behavior is consistent with a prior over values guiding efficiently coded value representations [37], the evidence remains correlational because values were not directly observed or manipulated and had to be inferred from behavior [45, 46]. Conversely, studies that have manipulated contextual value statistics have typically done so by altering the perceptual inputs from which values are constructed, making it difficult to determine whether the resulting effects arise from efficient coding in perception, in valuation, or in the transformations linking them. Because of this coupling, previous approaches could not causally isolate efficient coding across the coupled representational stages, leaving unclear whether behavioral biases reflect misperception, misvaluation, or both.

**Figure 1:**
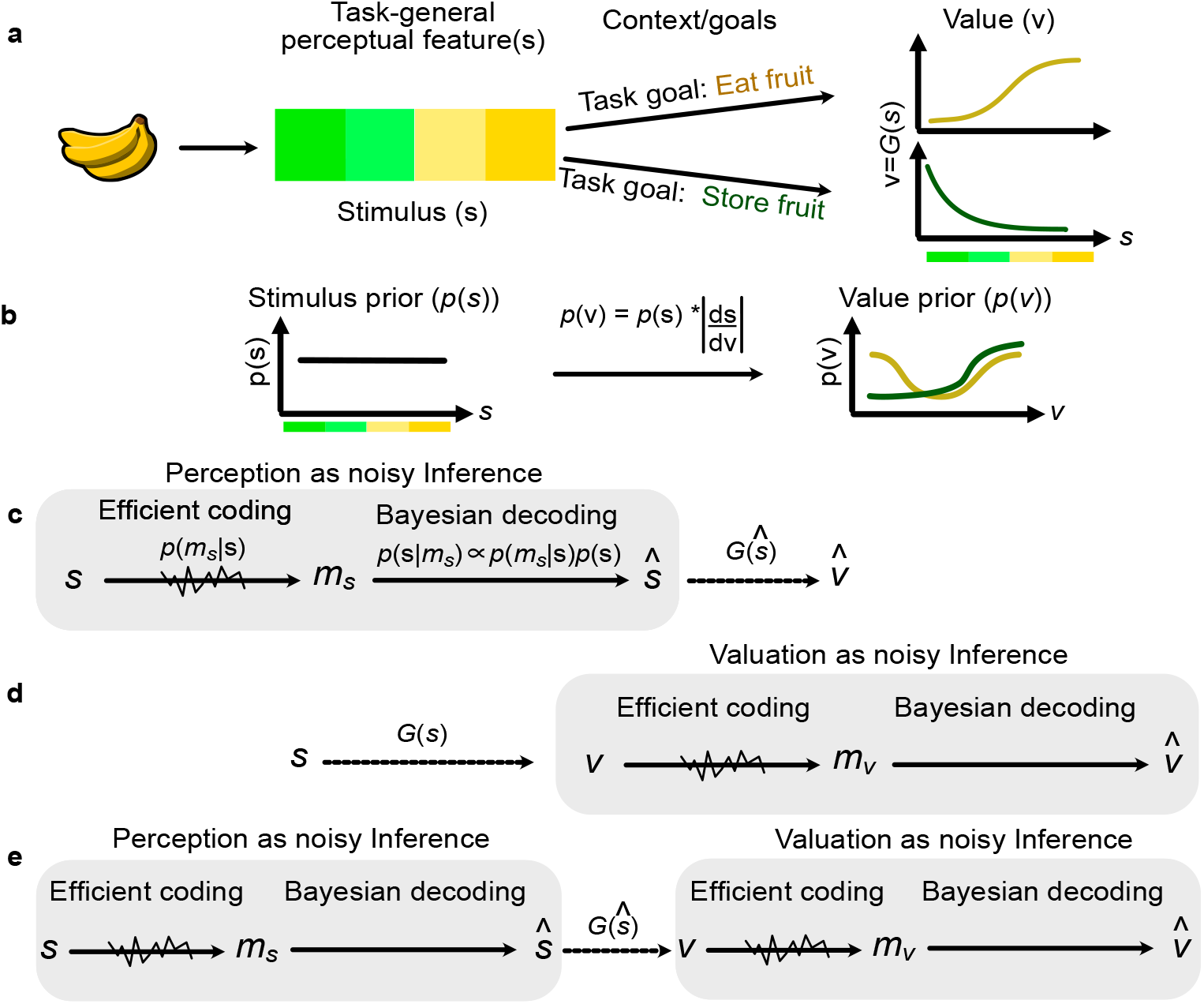
Competing architectures of efficient multi-stage representations for flexible behavior. **Fig. 1** (**a**) A single stimulus gives rise to stable, task-general perceptual features (here illustrated as hue, ranging from green to yellow) that are independent of goals. The same perceptual feature can be mapped onto different task-specific values depending on context or goals. For example, increasing greenness may decrease value when the goal is to immediately eat a banana, but increase value when the goal is to store it for later consumption. Thus, value is not an intrinsic property of the stimulus, but of the stimulus under a goal, motivating a distinction between perceptual and value representations. mapping *G*. Formally, 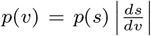 with *v* = *G*(*s*), where ^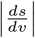^ is the absolute inverse slope of (**b**) With the stimulus distribution fixed, the value prior is determined solely by the stimulus-value *G*. Under a uniform stimulus distribution, shallower regions of the mapping yield higher value-prior density and steeper regions yield lower density. Different goals therefore induce distinct mappings *G*, reshaping the value distribution while leaving perceptual input statistics unchanged. (**c**) *Perception-only model*, in which efficient coding and Bayesian decoding operate exclusively in perceptual space. A stimulus feature *s* is efficiently encoded into a noisy representation *m*_*s*_ and Bayesian optimally decoded to get the mean percept *ŝ*, from which value is read out veridically, so behavioral biases arise solely from perceptual distortions. (**d**) *Value-only model*, in which perception is veridical and structure-dependent distortions arise in value space. Values *v* are efficiently encoded into a noisy representation *m*_*v*_ and Bayesian optimally decoded to get the mean inferred value 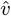 under context-dependent value structure. (**e**) *Multi-stage model*, in which perceptual and value representations occupy distinct spaces. A stimulus *s* is efficiently encoded into *m*_*s*_ and Bayesian optimally decoded to get the mean percept *ŝ*; the percept is mapped to a value *v* = *G*(*ŝ*), which is efficiently encoded into *m*_*v*_ and Bayesian optimally decoded to the mean inferred value 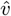. Efficient coding and Bayesian decoding operate independently at both stages, so statistical structure at each stage shapes inference. Behavioral output reflects the combined influence of perceptual and value distortions, which reinforce or compete depending on how structure aligns across stages.

The same coupling between perceptual and value distributions that makes multistage efficient coding difficult to disentangle experimentally also creates an opportunity. Because values are constructed through mappings from perceptual features, altering the stimulus–value mapping (for example by changing goals) can reshape value distributions even when perceptual input statistics remain unchanged. For instance, when the goal is immediate consumption, a supermarket containing mostly yellow ripe bananas will contain many high-value items; under a different goal, such as storing fruit for later consumption, the same perceptual environment will instead contain many low-value items. Thus, although value statistics cannot be manipulated independently of perceptual statistics under a fixed mapping or goal, changing the mapping itself, via the link between a given sensory feature and different goals, can alter value priors while leaving perceptual statistics fixed (Fig. 1b). This distinction also suggests that efficient coding may operate over different timescales across representational stages: Normative theories predict that adaptation should track the volatility of the represented distribution [47–49]. Perceptual features such as orientation, which follow relatively stable environmental statistics, exhibit efficient codes aligned with long-term priors [28, 29, 50]. In contrast, when stimulus statistics change rapidly across contexts, such as for contrast or luminance, sensory systems exhibit adaptive efficient coding that tracks recent stimulus distributions [51–53]. Because values are induced by goal-dependent stimulus–value mappings, their statistical structure is inherently context-dependent and should therefore show adaptive efficient coding over fast timescales. Here, we leverage this property to test whether behavior reflects adaptive efficient coding of values even when they are associated with relatively stable perceptual features, by manipulating value distributions independently of perceptual inputs through changes in stimulus-value mappings.

The ability to alter value distributions while holding perceptual input statistics fixed gives us a way to dissociate where along the perception–action processing pathway efficient coding operates and over what timescales it adapts. Specifically, we exploit this property to address three related questions: (1) How does noise at one representational stage propagate through multi-stage transformations to produce behavioral distortions? (2) Does efficient coding operate within a single representational stage or across multiple linked stages that jointly shape behavior? (3) Are the timescales of efficient coding different across stages, reflecting the distinct statistical stability of the distributions of variables they encode? To address these questions, we developed a computational modelling framework and experimental design that make these models qualitatively distinguishable. Specifically, we formalized three competing representational architectures in which efficient coding shapes only perceptual representations, only value representations, or both. We then exploited changes in the stimulus-value mapping while holding perceptual statistics constant, using orientation as our perceptual feature given its well-characterized long-term environmental statistics [29, 54]. This design generated distinct qualitative predictions across architectures and timescales, which we evaluated using preregistered statistical tests and formal model comparison. To preview, our results support a multi-stage representational architecture in which efficient coding operates independently at both perceptual and valuation stages. Nonlinear transformations between stages further reshape the propagated noise to the next stage, producing additional systematic biases and variability patterns in behavior. Furthermore, our results show that perceptual representations remain anchored in long-term environmental structure, whereas value representations adapt rapidly to short-term contextual priors.

## 2 Results

### Structure-driven efficient coding from perception to action

To dissociate where along the perception–action pathway efficient coding operates and over what timescales it adapts, we formalize three competing representational models that differ in where prior structure shapes inference through efficient coding and Bayesian decoding, and thereby in the systematic behavioral biases they predict (Fig. 1c–e; see Methods, Computational modelling, for full model specification and equations). All three exploit the fact that perceptual and value representations encode variables with different statistical stability: perceptual features such as orientation follow stable long-term environmental statistics, whereas values are induced by goal-dependent stimulus–value mappings and therefore vary rapidly with context (Fig. 1b). Formally, the value prior induced by a mapping *G* is 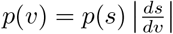, where 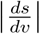 is the absolute value of the reciprocal of the local slope of *G* (defined for strictly monotonic mappings): shallow regions of the mapping compress many stimuli into few values, yielding higher value prior density, whereas steep regions spread stimuli more sparsely, yielding lower prior density. This makes it possible for efficient coding to operate at one stage but not the other, or at both with different adaptation timescales.

The first two models posit that efficient coding and Bayesian decoding exclusively operate within a single representational stage. Thus, under the *efficient-perception* model (model 1; Fig. 1c), efficient coding and Bayesian decoding act exclusively on task-general perceptual representations: a stimulus feature *s* is encoded into a noisy internal measurement *m*_*s*_ and decoded to yield a perceptual estimate *ŝ*(*m*_*s*_). Valuation then proceeds via a veridical transformation through the task-defined mapping *v* = *G*(*ŝ*), so that behavioral biases reflect stimulus statistics alone. In contrast, under the *efficient-valuation* model (model 2; Fig. 1d), perceptual representations are assumed to be veridical, and efficient coding and Bayesian decoding only operate at the valuation stage. Here, behavioral biases are shaped exclusively by the statistical structure of value space, not by stimulus statistics.

The third model posits that the same principles of efficient coding and Bayesian decoding operate across both stages, with independent statistical structure shaping inference at each (model 3; Fig. 1e). Under this *multi-stage efficient representation* model, the distribution of stimuli shapes efficient perceptual representations and the resulting distortions in perceptual space, while the distribution of values induced by the stimulus–value mapping additionally shapes distortions in value space. Because these sources of structure need not align, perceptual and valuation biases can either *reinforce* or *compete* with one another depending on task context, and behavioral output reflects the combined influence of multiple representational stages rather than a single locus of distortion. Crucially, this model uniquely predicts that behavioral biases due to perceptual efficient coding can persist, amplify, or reverse depending on whether perceptual and value priors align or conflict. Together, the *efficient-perception, efficient-valuation*, and *multi-stage efficient representation* models provide a principled framework for testing whether structure-driven biases arise in perceptual representations, value representations, or both, and for examining how shared computational principles are expressed across representations with distinct functional roles.

### Experimental dissociation of multiple representational priors and timescales

To test whether perceptual and value representations are governed by shared efficient coding principles operating separately on both types of representations, and over different timescales, we dissociated three sources of statistical structure: (i) long-term perceptual priors over stimuli, reflecting stable environmental regularities; (ii) shortterm perceptual priors imposed by uniform experimental sampling; and (iii) short-term value priors induced by condition-specific stimulus-value mappings (i.e., *G*(*s*)). The first two were held fixed and identical across conditions, while only the value prior was manipulated, allowing us to test both whether efficient coding operates across multiple representational stages that shape behavior, and whether the priors governing perception and valuation differ in their effective timescales.

To identify whether and how the long-term and short-term perceptual priors constrain the perceptual efficient coding, we focused on a stimulus dimension with well-characterized long-term environmental statistics: visual orientation. Cardinal orientations are overrepresented in natural scenes, and this long-term structure is known to shape perceptual representations through efficient coding [54]. In our experiment, orientations were sampled uniformly for all participants, across all phases and in all conditions, thereby imposing a uniform short-term perceptual prior that differs from the known long-term orientation prior (Fig. 3a). This allows us to dissociate long-term perceptual priors from short-term priors, and to test whether low-level perceptual representations continue to reflect stable environmental structure, consistent with prior work [27, 54]. Importantly, both long-term and short-term perceptual priors were identical across participants and conditions in our experiments, thus fixing the perceptual input statistics.

Unlike perceptual distributions, which reflect properties of the external world, value distributions are not intrinsic to the stimulus but are induced by mapping perceptual features onto task-defined outcomes. Because values are derived rather than directly sampled, the short-term prior over values is jointly determined by the short-term perceptual prior and the stimulus-value mapping (Fig. 1b): formally, 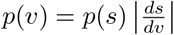 (see Fig. 1b and accompanying text for intuition). This coupling typically makes it difficult to isolate value priors experimentally. However, when perceptual inputs are sampled uniformly and held fixed, the stimulus–value mapping alone determines the resulting value distribution. We exploit this property by manipulating the orientation-value mapping while leaving perceptual input statistics unchanged, thereby selectively controlling short-term value priors without altering either long-term or short-term perceptual priors.

A total of 123 participants completed the task (Fig. 2a–c), in which participants first learned to associate visual stimuli with monetary values, analogous to how people learn that sensory features of everyday items, such as foods or clothes, predict their value. This orientation–value mapping was taught in two successive learning phases before valuation was measured in a no-feedback bidding task. After applying preregistered exclusion criteria, the final analyzed sample comprised 119 participants (mean age = 23.29 years; 54 female). In the first learning phase (*self-paced observational learning* ; Fig. 2a), we introduced participants to the full orientation–value mapping under low perceptual uncertainty (100% contrast). We presented high-contrast Gabor stimuli together with their associated values at 23 discrete orientations, spanning 7.5° to 172.5° in steps of 7.5°. Participants could freely move forward and backward through all stimulus–value pairings with unlimited viewing time to learn the mapping. In the second learning phase (*supervised training with feedback* ; Fig. 2b), participants actively reported values for the same 23 orientations under 100% contrast and received trial-by-trial feedback within a 10 s response window. We repeated each orientation 10 times in randomized order (230 trials total), ensuring equal exposure to every orientation– value pair during active learning. Thus, participants received feedback-driven practice with the same value-report format later used in the valuation task, but under easier conditions with higher stimulus contrast, a longer response window, and trial-by-trial feedback. Because feedback was provided only after each active value report, responses during the second learning phase could still reflect perceptual or valuation biases; the purpose of this phase was to provide additional supervised opportunity to learn the mapping before the no-feedback valuation task.

**Figure 2:**
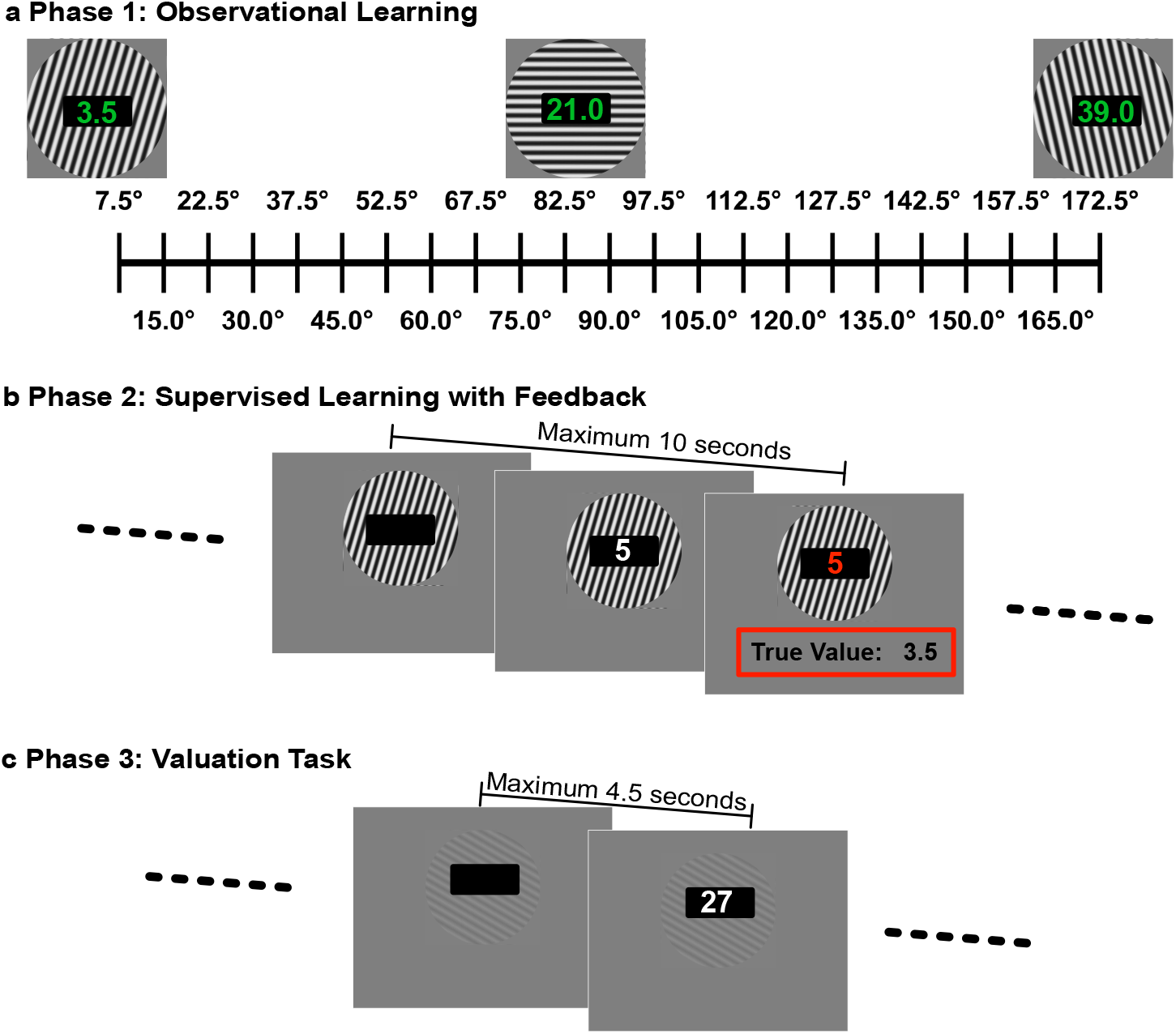
Three phase experimental paradigm. Fig. 2. Experimental paradigm and task structure. (**a**) Self-paced observational learning. Participants freely explored the full orientation–value mapping under low perceptual noise (100% contrast; 23 stimulus–value pairings, unlimited viewing time, self-paced). (**b**) Learning with feedback. Participants reported values for each orientation using a keypad and received trial-by-trial feedback (maximum response time: 10 s; 230 trials). (**c**) Valuation task. Orientations were presented at low contrast (2%) to increase perceptual uncertainty. Participants submitted monetary bids without feed-back within a 4.5 s response window (204 trials). Bids were elicited using a Becker–DeGroot–Marschak auction: on each trial, the bid was compared with a random price, determining whether the participant purchased the stimulus and the resulting trial payoff. Thus, bids had real financial consequences and measured participants’ valuation of the item; only these choices were used for hypothesis testing.

**Figure 3:**
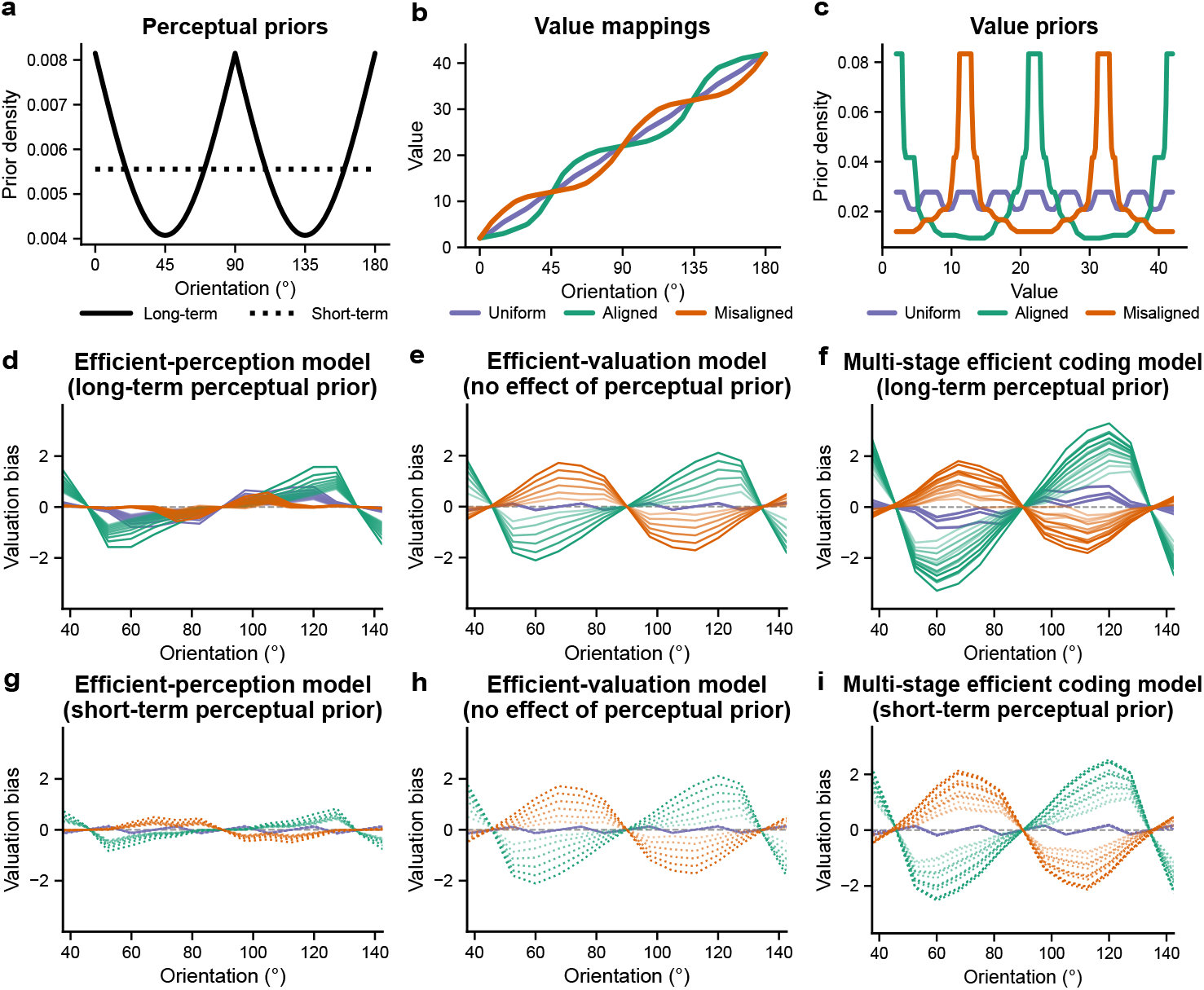
Predictions of perception, valuation and multi-stage efficient coding at different timescales. **Fig. 3** Throughout all figures, mapping conditions are color-coded consistently: *Uniform* = *purple, Aligned* = *green, Misaligned* = *orange*. Simulations implement efficient coding as a cumulative distribution function (CDF) transformation of representational space [27–29] followed by Bayesian decoding under explicit loss functions. (**a**) Perceptual priors over orientation: a long-term environmental prior reflecting natural orientation statistics (solid) and the uniform short-term perceptual prior imposed by the experimental stimulus distribution (dashed). (**b**) Orientation-value mappings defining the three experimental conditions (Uniform, Aligned, Misaligned). (**c**) Short-term value priors induced by each mapping under uniform orientation sampling. (**d-f**) Predictions when perceptual encoding is governed by the long-term environmental prior. (**d**) *Efficient-perception model* : orientation is efficiently encoded under the long-term prior with von Mises sensory noise (*κ*) and decoded under the long-term prior with a circular loss exponent *p*_per_ = 8 [55]; valuation is a deterministic readout, yielding biases that remain qualitatively similar in direction across mappings (with only minor mapping-dependent deviations due to the nonlinear stimulus-value transformation – Jacobian effects). (**e**) *Efficient-valuation model* : orientation perception lacks structure-dependent efficient coding distortions; values are efficiently encoded under the induced short-term value prior with Gaussian noise (*σ*) and decoded under squared-error loss (*p*_val_ = 2), producing no systematic bias under the Uniform mapping and opposite distortions under the Aligned and Misaligned mappings. (**f**) *Multi-stage efficient-coding model* : perceptual and value representations are independently shaped by their respective priors, yielding persistent bias under the Uniform mapping, amplification when perceptual and value priors align (Aligned), and a sign reversal with attenuation when they conflict (Mis-aligned). (**g-i**) Predictions when perceptual encoding is governed by the *short-term uniform prior*. (**g**) *Efficient-perception model* : predictions collapse toward minimal bias due to the absence of perceptual structure. (**h**) *Efficient-valuation model* : predictions are unchanged because distortions arise in value space. (**i**) *Multi-stage model* : predictions collapse toward valuation-only behavior, eliminating persistent bias under Uniform and restoring symmetry across Aligned and Misaligned mappings. Multiple curves within each prediction panel show model predictions across diagnostic ranges of the relevant noise parameters. In efficient perceptual model simulations, curves vary perceptual precision *κ*; in efficient valuation model simulations, curves vary value noise *σ*; and in multi-stage model simula-tions, predictions are shown across combinations of *κ* and *σ*. These parameter variations demonstrate that the qualitative signatures used for preregistered model discrimination are robust across plausible noise levels.

The valuation task (Fig. 2c), conducted after the learning phases described above, directly measured the subjective value that participants assigned to each stimulus using an incentive-compatible elicitation procedure standard in economic decision-making. On each trial, participants placed a bid for the presented Gabor under a Becker–DeGroot–Marschak (BDM) auction mechanism. Each trial had real financial consequences: participants received a trial budget of 42 CHF, submitted a bid, and this bid was compared with a randomly drawn price. If the bid was at least as high as the random price, the participant purchased the stimulus at that price and earned the remaining budget plus the stimulus value; if the bid was lower, no purchase occurred and the participant retained the full trial budget. Trial earnings were summed across the valuation task and converted into real payment at a fixed rate, in addition to the show-up fee. Under this mechanism, participants maximize their expected payoff by bidding the stimulus’s true subjective value, so bids provide a direct, trial-by-trial readout of their value estimates. To increase perceptual uncertainty, we presented Gabor stimuli at low contrast (2%), withheld feedback, and gave participants a 4.5 s response window. We focused testing on a preregistered window of interest comprising 17 orientations spanning 30° to 150° in steps of 7.5°, each repeated 12 times (204 trials total). We chose this restricted testing window because it is the region where the predictions of the three representational models diverge most strongly, so that data collected here were maximally diagnostic for discriminating between them.

To dissociate efficient coding at the perceptual and valuation stages, we needed to manipulate the value prior independently of the perceptual prior. We achieved this by randomly assigning participants to one of three preregistered orientation–value mappings (Fig. 3b; Supplementary Table 1), each of which induced a distinct short-term value prior under identical perceptual input statistics (Fig. 3c). In the *Uniform* mapping, values increased linearly with orientation, producing an approximately uniform value prior; because value was a linearly scaled readout of orientation, this condition provided a simple baseline for orientation-linked biases expressed through value reports. In the *Aligned* mapping, we mapped orientations near the cardinal axes onto a disproportionately dense region of value space, inducing a value prior peaked at values corresponding to cardinal orientations and thus aligned with long-term perceptual structure. In the *Misaligned* mapping, we inverted this relationship, producing a value prior peaked at values corresponding to oblique orientations and thus misaligned with the long-term prior distribution of those orientations. Across all three conditions, we held perceptual priors fixed and identical, while value priors differed systematically in their alignment with long-term perceptual structure. This manipulation isolated short-term value priors and probed valuation as a fast, context-dependent representational stage: because changing the mapping (and thus the implicit goal) reshapes the value distribution while leaving perceptual inputs untouched, we can attribute any resulting behavioral differences across conditions specifically to value-stage coding.

Because extensive work shows that orientation perception reflects stable naturalistic priors [27, 54], we expect perceptual encoding to remain governed by long-term environmental structure even though the experiment imposes a uniform short-term orientation distribution. Under this regime, the three representational models make sharply divergent qualitative predictions for the bias patterns across the three mappings (Fig. 3d–f), and these signatures remain stable across the diagnostic ranges of the relevant noise parameters shown by the multiple curves in Fig. 3. A caveat is that biases can also arise from the nonlinear transformation between perceptual and value representations: when noisy perceptual estimates pass through a nonlinear stimulus– value mapping, the transformation reshapes the geometry of perceptual noise and can produce systematic biases even without efficient coding (Jacobian effects). Our simulations explicitly include these transformation-induced biases, so the remaining qualitative differences isolate efficient coding at one or more representational stages.

Each column of Fig. 3d–f shows the bias pattern predicted by a model in which efficient coding operates at a different locus: perception (d), valuation (e), or both (f). Under the *efficient-perception model* (Fig. 3d), biases are exclusively governed by longterm perceptual priors and therefore remain qualitatively similar across mappings: changing the stimulus–value mapping cannot reverse the sign of the bias pattern, and the only mapping-dependent differences are the small Jacobian effects described above. By contrast, under the *efficient-valuation model* (Fig. 3e) biases are exclusively governed by the induced short-term value prior. Note that under the efficient valuation model, the *Uniform mapping* shows no systematic bias, whereas the Aligned and Misaligned mappings produce symmetric but opposite distortions, yielding a complete reversal in bias direction between these conditions. A robust sign change of the observed biases between Aligned and Misaligned would therefore provide a clear qualitative signature of efficient coding in value space, driven by fast adaptation to short-term contextual priors, over and above any perceptual contribution. Finally, the *multi-stage efficient-coding model* predicts a combination of these two signatures (Fig. 3f): the Uniform mapping retains a perceptual-like repulsive bias around the cardinal, the Aligned mapping shows amplified repulsion because perceptual and value priors reinforce one another, and the Misaligned mapping shows a sign-reversed but attenuated bias because the two priors compete (Fig. 3f). Thus, short-term value priors still drive a reversal between Aligned and Misaligned, but the overall pattern becomes asymmetric: biases are amplified when perceptual and value priors reinforce one another (Aligned) and reversed but attenuated when they compete (Misaligned), while perceptual-like biases persist under the Uniform mapping. Thus, the Uniform mapping isolates perceptual distortions without value-stage contributions, because under this mapping values are simply linearly scaled orientations. The Aligned–Misaligned comparison tests whether value-stage efficient coding reinforces or opposes this baseline. Together, this asymmetric combination across the three mappings both identifies multi-stage efficient coding and reveals that its two stages operate at distinct timescales — fast contextual adaptation in valuation, and slower environmentally anchored structure in perception.

Although orientation biases are thought to reflect stable long-term environmental statistics rather than short-term adaptation [27, 54], for completeness we also simulate the alternative case, in which perceptual encoding adapts fully to the experimentally imposed short-term uniform orientation prior (Fig. 3g–i). In this limiting case, the efficient-perception model predicts near-zero biases apart from small Jacobian effects, because the perceptual prior is uniform. The multi-stage model also collapses toward the efficient-valuation pattern, because the perceptual stage no longer contributes structured distortions. Thus, under a short-term uniform perceptual prior, value-only and multi-stage models are not qualitatively separable from the bias patterns alone.

Comparing these two regimes also clarifies the diagnostic logic of the experiment. Across both short- and long-term perceptual-priors, an Aligned–Misaligned reversal indicates efficient coding in value space driven by fast adaptation to short-term value priors. What uniquely identifies the multi-stage architecture is the additional presence of a residual, perceptually driven bias under Uniform, together with asymmetric amplification versus attenuation across Aligned and Misaligned. Observing this full asymmetric pattern would show that behavior reflects both efficient coding in valuation at a fast contextual timescale and efficient coding in perception at a slower environmental timescale. Together, these preregistered qualitative signatures provide a decisive test of whether behavior reflects (i) perceptual distortions driven by priors at slower environmental timescales, (ii) multi-stage efficient coding in perception and valuation, and (iii) value-based distortions driven by fast short-term priors.

### Behavioral biases reveal multi-stage efficient coding with fast contextual value priors and slow long-term perceptual priors

We evaluated the behavioral evidence for multi-stage efficient coding in three complementary ways. First, we used a qualitative, model-guided inspection of the predicted and observed bias functions to assess whether the empirical pattern matched the preregistered diagnostic signatures of the competing architectures (OSF preregistration: https://osf.io/fcnam/overview). Second, we tested these signatures using preregistered model-free mixed-effects analyses of valuation bias around the cardinal orientation. Third, we compared the full computational models by fitting them to trial-wise bids and evaluating their relative likelihood. This structure allowed us to ask whether the same diagnostic pattern was visible qualitatively, supported statistically with preregistered model-free analyses, and favored by formal model comparison.

In the qualitative inspection, changing only the stimulus–value mapping, and thereby selectively reshaping the short-term prior in value space while holding perceptual input statistics fixed, was associated with pronounced mapping-dependent differences in the direction and magnitude of valuation biases as predicted in our pre-registration (Fig. 4d). Most tellingly, biases around the cardinal orientations reversed sign between the Aligned and Misaligned conditions despite identical perceptual inputs. This sign reversal is incompatible with an efficient-perception model, in which distortions are fixed in stimulus space and therefore retain the same direction across mappings (Fig. 4a); instead, it provides a direct signature of fast and flexible, adaptive efficient coding in value space, with short-term value priors rapidly reconfiguring value-space distortions. The efficient-valuation model, however, could not fully capture the data either (Fig. 4b). Bias magnitudes were asymmetric, with stronger repulsion when perceptual and value priors reinforced one another (Aligned) than when they opposed one another (Misaligned), and systematic repulsive structure persisted even under the Uniform mapping. These features are consistent with a slower perceptual contribution anchored in long-term orientation structure that interacts with the fast value-stage distortions. Together, the combination of (i) mapping-dependent sign reversal, (ii) persistence under Uniform, and (iii) asymmetric amplification versus attenuation matches the diagnostic triplet uniquely predicted by the multi-stage efficient-coding model, in which fast value priors operate alongside a slower perceptual component (Fig. 4c; Fig. 3). Next, we tested these qualitative signatures statistically.

**Figure 4:**
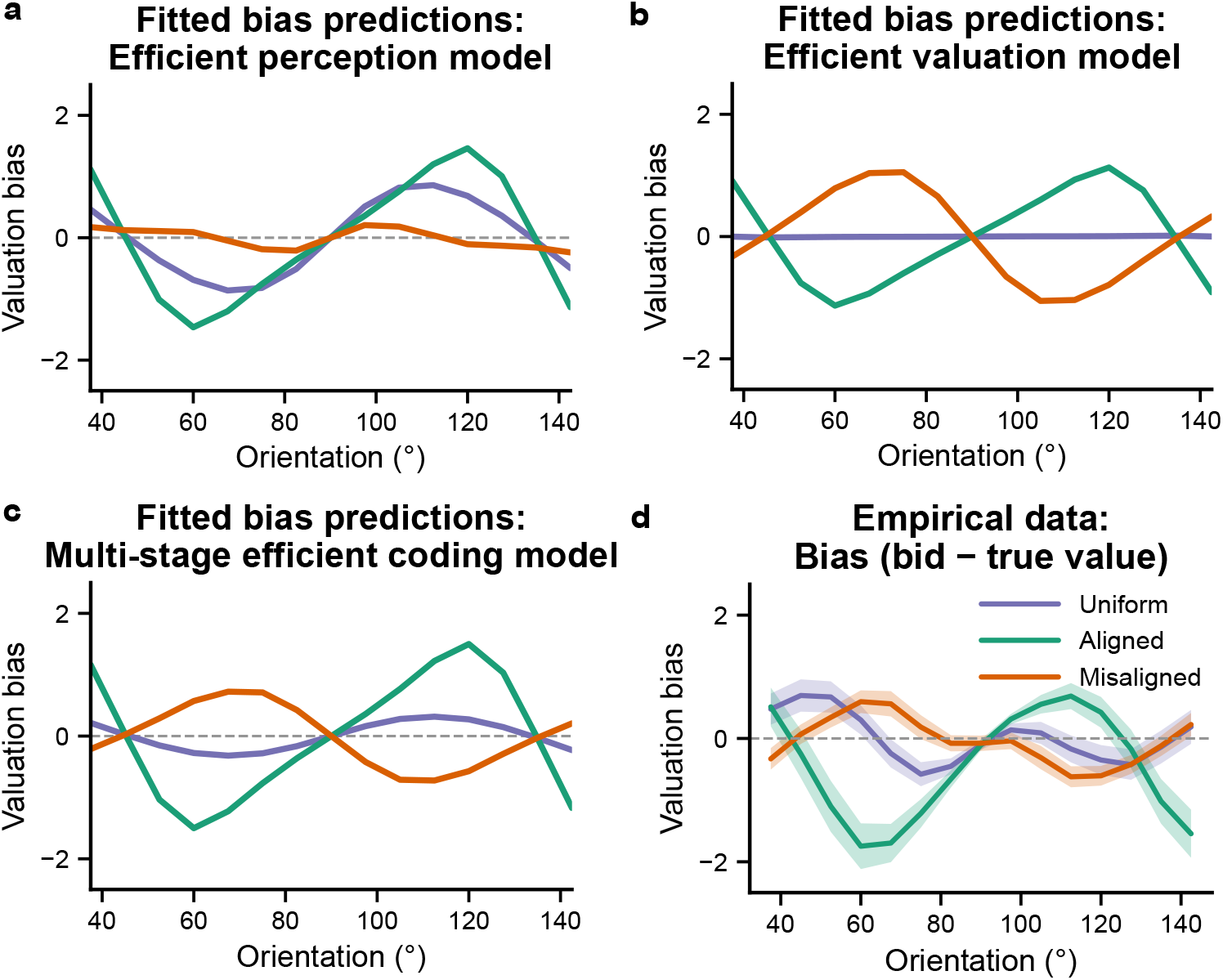
Empirical data supports multi-stage efficient coding at different timescales. **Fig. 4** Panels show best-fitting model predictions (left columns; see methods for details) and empirical data for valuation bias (Phase 3), focusing on the cardinal region around 90°. For all model fits shown here, perceptual encoding is assumed to be governed by the long-term environmental prior over orientation (slow perceptual timescale), while short-term contextual structure is manipulated in value space via the stimulus–value mapping (see Fig. 3). (**a**) Best-fitting predictions of the *efficient-perception* model, in which efficient coding and Bayesian decoding operate only in perceptual space and valuation is a deterministic readout of perceptual estimates. This model predicts qualitatively similar bias patterns across mappings, with direction primarily determined by perceptual structure. (**b**) Best-fitting predictions of the *efficient-valuation* model, in which perceptual representations lack structure-driven efficient coding and efficient coding operates only in value space. This model predicts no systematic bias under the Uniform mapping and opposite distortions under the Aligned and Misaligned mappings, reflecting short-term value priors. (**c**) Best-fitting predictions of the *multi-stage efficient-coding* model, in which perceptual and value representations are independently shaped by their respective priors. This model predicts a diagnostic triplet pattern: persistent bias under the Uniform mapping, amplification when perceptual and value priors align (Aligned), and a sign reversal with attenuation when they conflict (Misaligned). (**d**) Empirical valuation biases, computed as bias = bid *−* true value (positive values indicate overestimation). Data exhibit the predicted triplet signature across mappings, consistent with multi-stage efficient coding and rapid, context-dependent distortions induced by short-term value priors. Curves are shown as a function of stimulus orientation and averaged across participants within each mapping condition. Alternative fits under the assumption that perceptual encoding is governed by the short-term uniform prior yield qualitatively value-like predictions (cf. Fig. 3g–i).

In the preregistered model-free analyses, we tested these qualitative signatures statistically using the observed responses in a fixed window around the cardinal axis (65°–115°). We had preregistered four hypotheses: mapping-dependent differences in bias slope (H1), persistence of repulsive bias under the Uniform mapping (H2), amplification of repulsive bias in the Aligned condition relative to the Uniform condition (H3), and a sign reversal of bias slope within the Misaligned condition (H4). We defined valuation bias as bias = bid *−* true value and treated orientation as a continuous predictor centered at 90° (*θ*_*c*_ = *θ* − 90°). This preregistered window was chosen to provide a simple local linear summary of bias around the cardinal axis, where the competing models make clearly separable predictions, while avoiding value-mapping boundaries that could introduce additional boundary-related biases across all mappings [42, 56]. We note, however, that both model predictions and empirical bias functions diverge across a broader, nonlinear orientation range (Fig. 4d). The central prediction of the multi-stage model is that changing only the stimulus-value mapping should rapidly reshape how bias varies with orientation, despite identical perceptual input statistics. Consistent with this, the preregistered mixed-effects model (bias ∼ *θ*_*c*_ *×* mapping + (1 | Participant); 9,615 observations, 119 participants) revealed strong mapping-dependent differences in bias slope (H1). Relative to Uniform, the Aligned mapping produced a substantially steeper positive slope 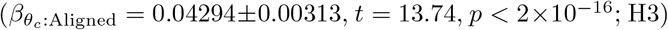, whereas Misaligned produced a strongly reduced and sign-opposed slope 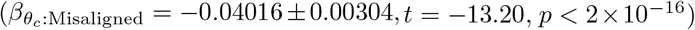. The strong difference in bias patterns between the Aligned and Misaligned conditions (0.08310 ± 0.00306, *t*(9609) = 27.12, *p* = 3.64 *×* 10^−156^) quantitatively confirms the qualitative reversal in Fig. 4d. This mapping-induced inversion shows that the canonical repulsive bias around the cardinal orientations can be reshaped, and even reversed, by rapid contextual structure in value space despite identical stimulus statistics, a result incompatible with purely perceptual accounts.

Having shown that value-space structure can reshape and even reverse these biases, we next asked whether perceptual inference shaped by long-term orientation structure contributes in parallel. Two features of the behavior pointed to this parallel perceptual contribution. First, under the Uniform mapping, where the induced short-term value prior is flat, biases nevertheless showed reliable repulsion around the cardinal (H2), reflected in a significantly positive slope 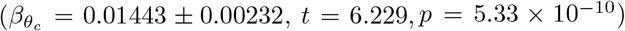. Second, bias magnitudes were systematically asymmetric across the two non-uniform mappings, with the absolute slope being substantially larger under Aligned than Misaligned (H3 above), consistent with reinforcement of orientation biases when perceptual and value priors align and attenuation when they compete. This asymmetry is not expected under accounts in which perceptual encoding is governed by the uniform short-term stimulus statistics, which predict minimal bias under Uniform and a more symmetric Aligned–Misaligned pattern (Fig. 3g–i). Finally, the preregistered within-condition test confirmed the predicted slope reversal in Misaligned (H4): fitting bias ∼ *θ*_*c*_ + (1|Participant) within Misaligned trials only (42 participants) yielded a significantly negative slope 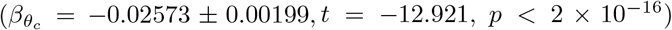. Together, these results reproduce the full pattern predicted by the multi-stage efficient-coding model, in which short-term value priors rapidly reconfigure valuation distortions while long-term perceptual structure shapes behavior in parallel. Full mixed-effects model results are reported in Supplementary Table 3.

In the formal model comparison, we then asked whether the same pattern was favored at the level of trial-wise bids by comparing six candidate models evaluating how well competing architectures explain the full distribution of participants’ bids. We compared six candidate models, formed by crossing three architectures (*Perceptual-only, Value-only*, and *Multi-stage*) with two assumptions about the timescale of the perceptual prior: perceptual encoding and decoding governed either by (a) the long-term environmental orientation prior or (b) the short-term uniform experimental prior. We fit all models at the individual-participant level to Phase 3 trial-wise valuation data using a fixed lapse rate of 0.01, an important safeguard against a few random or inattentive responses dominating the likelihood and distorting the fitted parameters (though the fits were highly robust to its exact value; Supplementary Table 2). Across both timescale configurations, the *Multi-stage* model provided the best account. Under long-term environmental priors, it achieved the lowest total NLL (64266.30) and was strongly favored by both AIC (129008.61) and BIC (129787.66), outperforming *Value-only* (AIC = 130569.03; BIC = 130958.56) and *Perceptual-only* (AIC = 133691.81; BIC = 134081.34). The same architecture was preferred when perceptual encoding instead reflected the short-term uniform prior (AIC = 129292.69; BIC = 130071.74). Within the *Multi-stage* architecture the best overall fit was obtained when perceptual representations remained anchored to long-term environmental structure while valuation representations were rapidly reshaped by short-term contextual priors. Full results across all architectures, perceptual prior assumptions, and lapse rates are reported in Supplementary Table 2; best-fitting parameter estimates (*κ*_*r*_ and *σ*_rep_) for all models and conditions are reported in Supplementary Table 4. Together with the preregistered bias signatures and model-free tests, these results provide three lines of evidence that adaptive valuation behavior reflects multi-stage perceptual and value representations under efficient coding at distinct timescales. Model recovery confirmed that the three mapping conditions jointly render the competing models identifiable at the group level (Supplementary Fig. 10).

### Behavioral variability patterns are consistent with multi-stage efficient coding

We next asked whether trial-to-trial behavioral variability shows signatures consistent with multi-stage efficient coding as well. The logic is simple: as noisy perceptual representations are transformed into value representations, efficient coding at each stage and the nonlinear stimulus–value mapping between them locally expand or compress internal noise. Behavior should therefore exhibit structured variability across stimulus space, with regions of expansion producing more variable bids and regions of compression producing less variable bids. We use these signatures not to adjudicate between models (the candidate architectures make similar variability predictions; model adjudication was carried out through the bias analyses above) but to ask whether the variability we observe is consistent with behavior guided by sequentially constructed, noise-transformed value representations.

Simulations of the best-fitting multi-stage model produce mapping-dependent variability profiles, with elevated variability where transformations expand representational space and reduced variability where they compress it (Fig. 5a). Empirical bid variability showed the same qualitative ordering across conditions (Fig. 5b, white shaded regions): highest in Aligned, lowest in Misaligned, and intermediate under Uniform. Away from the cardinal, predicted and observed variability profiles closely aligned, indicating that behavioral variability reflects structured transformations of internal noise rather than uniform response noise.

**Figure 5:**
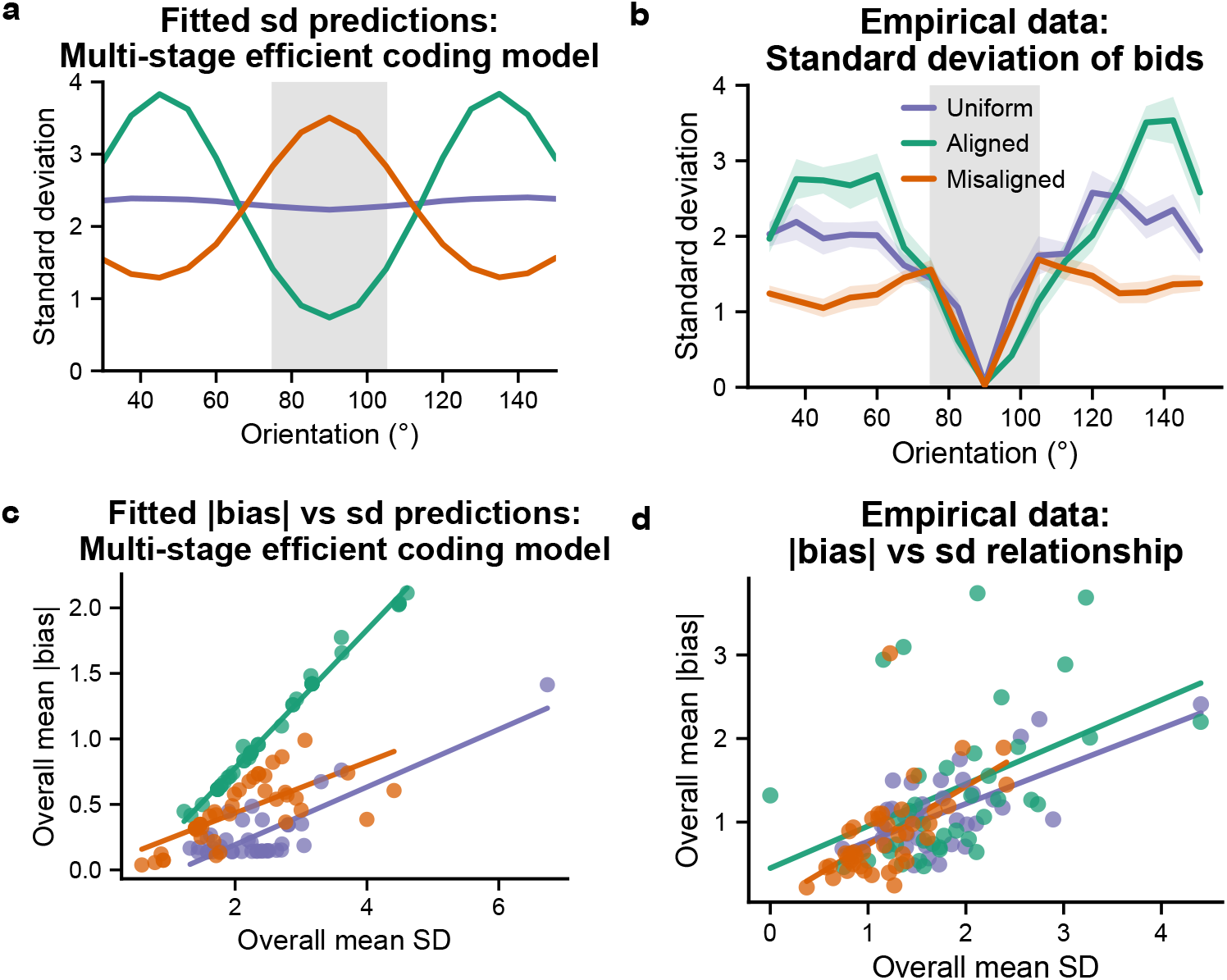
Multi-stage efficient coding model captures co-occuring variability patterns. **Fig. 5** (**a**) Predicted response variability (standard deviation of model-implied bid distribution per orientation) as a function of stimulus orientation, under the best-fitting multi-stage efficient-coding model in which perceptual and value representations are sequentially formed and shaped by prior structure at distinct timescales. The model predicts elevated variability in regions where multi-stage warping expands representational space and reduced variability where it compresses representations. (**b**) Empirical response variability in Phase 3, computed as the within-participant standard deviation of submitted bids at each tested orientation and then averaged across participants (shaded regions denote ± s.e.m.). Empirical variability exhibits the same qualitative ordering across mappings as predicted by the model, with highest variability in the Aligned condition, lowest variability in the Misaligned condition, and intermediate variability under the Uniform mapping. Deviations between model and data are concentrated near the 90° cardinal (grey shaded region), where empirical variability is markedly reduced. (**c**) Model-predicted relationship between overall distortion magnitude and overall response variability across participants, quantified as mean absolute bias (averaged across orientations) versus mean predicted bid standard deviation (averaged across orientations), shown separately for each mapping condition. (**d**) Empirical bias–variability relationship, showing that participants with larger systematic valuation biases also exhibit greater trial-to-trial variability (Uniform: *r*(37) = 0.657, *p <* .001; Aligned: *r*(36) = 0.440, *p* = .006; Misaligned: *r*(40) = 0.587, *p <* .001).

A striking exception emerged near the 90° cardinal (grey shaded region), where variability collapsed across all mapping conditions to near-deterministic levels. Because the multi-stage model accounts well for variability across the rest of stimulus space, this localized reduction is particularly informative: it reveals an additional stabilizing influence not captured by the continuous representational assumptions of the base model. We return to this anomaly in the next subsection.

Sequential noise propagation across multiple stages also predicts a within-subject relationship between bias and variability. If both arise from the same underlying representational noise transformed across stages, participants with noisier representations should show both larger systematic distortions and greater trial-to-trial variability. Indeed, participants with larger mean absolute valuation biases also showed greater response variability in all three mapping conditions (Uniform: *r*(37) = 0.657, *p <* .001; Aligned: *r*(36) = 0.440, *p* = .006; Misaligned: *r*(40) = 0.587, *p <* .001; Fig. 5c,d).

Together, these results show that sequential transformations of noisy internal representations not only shape systematic biases, but also behavioral variability across stimulus space. Furthermore, the localized variability collapse near the cardinal reveals a boundary condition of the continuous-inference account, which we examine next.

### Categorical stabilization at the horizontal cardinal

Although the multi-stage efficient-coding model guided by long-term perceptual priors and short-term valuation priors accounts for the dominant structure of valuation bias, response variability across orientation space, and the observed bias–variability coupling across subjects, it systematically underestimates a highly localized and mapping-invariant empirical signature near the 90° cardinal. Around the horizontal orientation, participants’ valuation responses exhibited an abrupt collapse of trial-to-trial variability, approaching deterministic behavior across all mapping conditions (Fig. 5b). The strong specificity of this deviation near 90° indicates that an additional stabilizing influence emerges selectively near the cardinal orientation for our task paradigm.

A key feature distinguishing our paradigm from classical perceptual matching experiments is that observers learned stable stimulus–value associations and reported learned numerical values, while orientations were sampled at a relatively coarse resolution (7.5° spacing). Under these conditions, the absence of variability at 90° suggests that internal precision for the learned value associated with the horizontal orientation exceeds the effective resolution of the task. Importantly, the perceptual prior (|2 − sin(*x*) | approximation) used in the model constitutes a smooth approximation, whereas environmental orientation statistics exhibit sharper peaks at cardinal orientations that cannot be fully captured by such parameterizations [54]. When perceptual precision locally exceeds the spacing between experimentally probed orientations, observers can reliably determine whether a stimulus lies below, at, or above the cardinal, eliminating category-level confusions. Behavior therefore appears effectively categorical at the resolution of the experiment, even though the underlying perceptual representations may remain continuous.

Because our experimental design does not independently manipulate stimulus spacing and response resolution, the mechanism giving rise to this regime cannot be identified here. Rather than modifying the perceptual prior or introducing additional perceptual mechanisms, we interpret this effect as reflecting that the smooth prior approximation underestimates local encoding precision from the long-term perceptual prior. We therefore model only the behavioral consequences of this mismatch using a minimal descriptive extension. The extension leaves perceptual and value inference unchanged but assumes that category membership relative to the cardinal (below, at, or above 90° at the experimental resolution) is inferred without error. Operationally, inferred value distributions are constrained to remain within the category implied by stimulus orientation, preventing category-crossing responses while preserving continuous estimation within categories. This implementation provides a mechanism-agnostic proxy for the behavioral regime in which increased precision removes category-level confusions without altering the underlying encoding or decoding processes.

As shown in Fig. 6, incorporating cardinal categorical stabilization closes the local gap between predicted and observed variability near the cardinal while leaving the broader predictions of the multi-stage model intact. Model predictions (Fig. 6a–c) demonstrate that bias profiles and bias–variance relationships remain unchanged, while variability near the cardinal is now accurately captured. Empirical data (Fig. 6d–f
) exhibit the same pattern, with the extended model reproducing both the global structure of behavior and the pronounced reduction in variability around 90°. Consistent with this qualitative improvement, the extended model provides a better account of trial-wise responses than the base multi-stage model (AIC = 123346.18; BIC = 124125.23; parameter estimates in Supplementary Table 4). Crucially, however, note that the important effects adjudicating between the different models remain unchanged in this model, so that our preregistered analyses are not affected by this additional effect and the way we model it.

**Figure 6:**
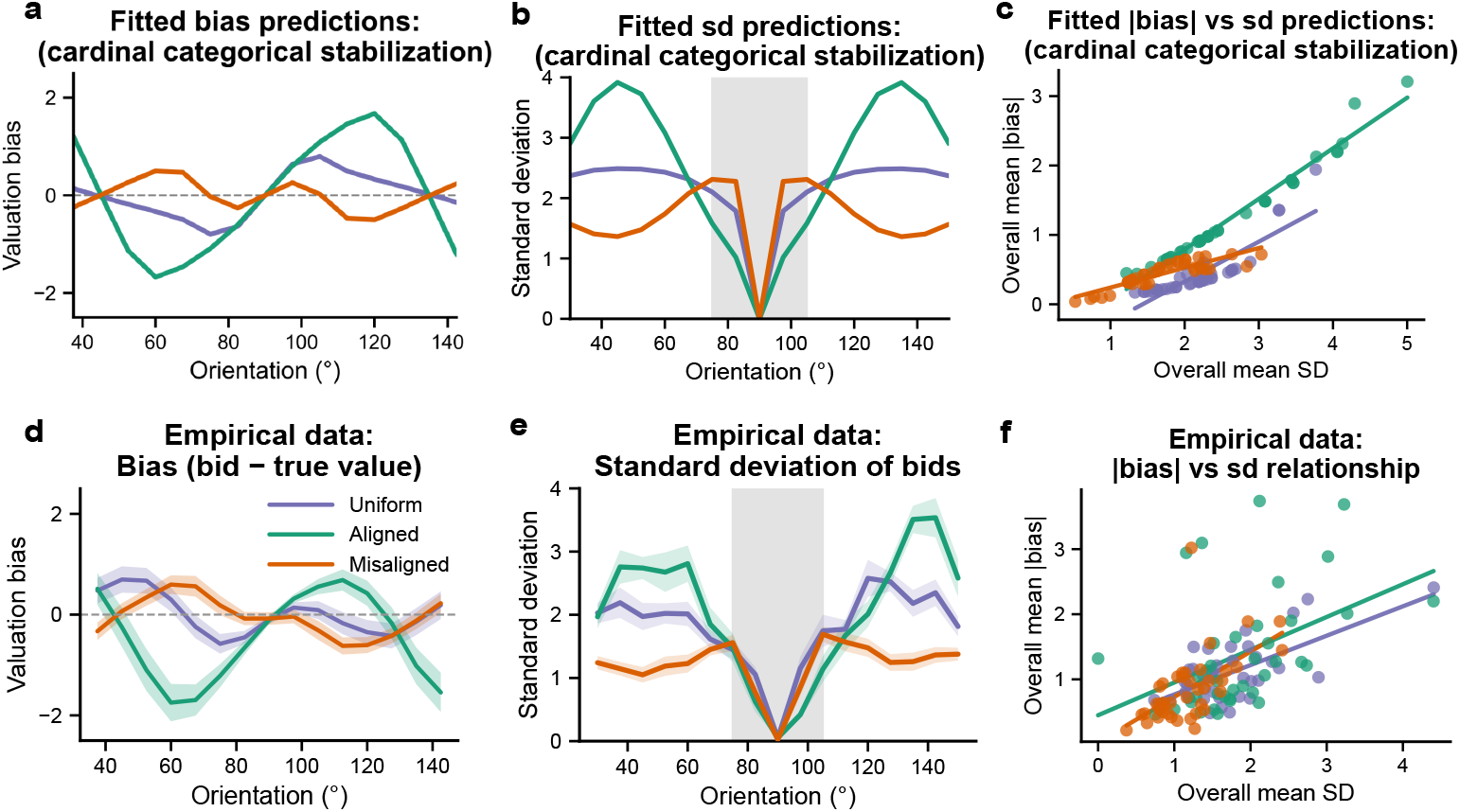
Additional categorical stabilization at cardinal explains deviations. **Fig. 6** Panels compare predictions of the multi-stage efficient-coding model extended with a descriptive cardinal categorical stabilization constraint (a–c) to empirical data (d–f). (**a**) Predicted valuation bias as a function of stimulus orientation. The extension leaves the bias structure largely unchanged relative to the base multi-stage model, preserving the mapping-dependent distortions explained by multi-stage efficient coding. (**b**) Predicted trial-to-trial response variability (standard deviation of bids per orientation). Incorporating cardinal categorical stabilization selectively eliminates the variance discrepancy near the 90° cardinal while maintaining variability structure elsewhere across orientation space. (**c**) Predicted relationship between absolute bias magnitude and response variability across participants, showing that the bias–variance coupling generated by multi-stage noisy representations is preserved. (**d–f**) Corresponding empirical measurements of bias, variability, and the bias–variance relationship. The extended model captures both the global behavioral signatures and the highly localized reduction in variability near the cardinal.

This descriptive extension was not preregistered and is not intended to adjudicate between representational architectures, which was addressed by the analyses above. Rather, it highlights an additional phenomenon revealed by our task: when internal representational precision exceeds the resolution afforded by the test values used in the experiment, localized categorical stabilization can emerge and strongly shape observable behavior. Determining when such precision-dependent effects arise, and how they depend on stimulus resolution, learning, and perceptual priors, represents an important direction for future work.

## 3 Discussion

A long tradition in the cognitive and neural sciences treats behavior as the output of hierarchically organized representations, from Marr’s account of vision as a sequence of constructed descriptions of the world [57] to evidence that cortical hierarchies progressively transform sensory inputs into increasingly abstract task-relevant variables [2, 16, 17, 58–61] that support behavior across domains [10, 11, 14, 15]. Yet characteristic patterns of behavior are typically modeled as arising from a single locus, usually related to perceptual inference or value / utility functions [3–5, 23–25, 27, 55]. However, this single-locus framing is challenged by work showing that choice variability can be dissociated into sensory, inferential, and response-selection components, with substantial suboptimality arising from limited precision in post-sensory inference rather than from sensory or response noise alone [62, 63]. In line with such considerations, our results argue that systematic behavioral patterns are best explained by sequentially formed, efficiently encoded representations that are Bayesian-decoded at multiple stages along the perception–action pathway, consistent with the broader view that behavior reflects efficient use of limited computational resources [64–66]. In doing so, our work directly addresses the recently highlighted question at which levels of the neural processing stream, and over which temporal scales, efficient adaptations should shape sensory information for behavior [44]. The mappings that link these stages also reshape the geometry of internal noise, producing additional distortions beyond those driven by efficient coding and Bayesian decoding within each stage. When distortions are attributed to a single locus, the resulting behavioral patterns can look inconsistent or puzzling; when the full sequence of representational transformations is considered, the same patterns fall out as predictable consequences of efficiency and optimal decoding operating across stages.

The domain of economic valuation and choice illustrates the point. A large literature documents that observed preferences vary systematically with context, framing, and elicitation procedure [67–70], which is difficult to reconcile with classical economic theories in which behavior reflects stable preferences captured by an *as if* utility function [3]. Recent work has begun to address parts of this puzzle by showing that distortions in economic choice can originate in perceptual representations, with efficient coding and Bayesian decoding shaping how sensory inputs feed into downstream valuation and choice [32, 39–42, 71, 72]. Our results extend this view in two ways. First, even when the stimulus–value mapping is deterministic, nonlinear transformations between representational stages can reshape the consequences of upstream noise: when noisy perceptual estimates are mapped into value space, local expansion or compression of the mapping changes how perceptual uncertainty propagates into valuation, producing Jacobian-related biases and variability differences over and above the biases generated by efficient coding within each stage. Second, value representations themselves can be governed by efficient coding, with priors that adapt rapidly to short-term context. Understanding behavior therefore requires attending to the full representational pathway: the constructed representations that directly guide action (here, value), the lower-level representations from which they are built (here, sensory features), and the deterministic mappings that link them. Efficient coding at each representational stage can introduce stage-specific distortions, while nonlinear mappings determine how upstream noise and distortions are propagated downstream. The framing extends naturally beyond the present task: whenever behavior depends on abstract constructed variables, the same multi-stage logic should apply.

A second implication of this study concerns the temporal scales over which efficient coding shapes representations across stages. In our task, perceptual distortions were best explained by long-term environmental statistics over orientations, whereas valuation distortions tracked short-term contextual priors induced by the stimulus–value mapping. This dissociation is consistent with normative accounts predicting that the timescale of efficient coding should reflect the statistical dynamics of the represented variable [47–49]. In sensory systems, efficient representations operate across multiple temporal regimes, from stable codes aligned with persistent environmental structure to adaptive responses that track recent stimulus statistics [28, 29, 50–53]. Our results suggest that the same principle extends beyond perception to constructed representations such as value, whose distributions are inherently shaped by changing task goals and contexts. In the present task, the relevant value prior was volatile by construction: changing the stimulus–value mapping immediately changed the distribution of values while leaving the perceptual input distribution fixed. Rapid value-stage adaptation is therefore consistent with the idea that representations should update quickly when the statistical structure they encode is unstable, but remain anchored when that structure is stable. This parallels work on adaptive learning in volatile environments, where learning rates are modulated by estimates of environmental change [73**?**, 74], and efficient-adaptation accounts, where coding resources should be recalibrated according to the stability of the current context [49**?**]. Although these literatures address different computational problems, they converge on a shared principle: both learning dynamics and representational structure should adjust across temporal scales according to the stability of the variables being inferred. This also suggests that not all value priors need to adapt equally rapidly. Values learned repeatedly across contexts, value distributions arising from stable goals, or values acquired early in development may be anchored by more stable statistical structure and may therefore update more slowly. Such persistent value priors could stabilize behavior in familiar environments, but may become maladaptive when they preserve outdated or negative expectations, linking the present framework to computational accounts in which psychiatric symptoms reflect rigid priors, biased belief updating, or impaired adaptation to changing environmental structure [75–77].

The present study examines how noise and resource limitations shape behavior once information is transformed across multiple stages, but it does not address why cognition may be organized this way. One plausible answer is that multi-stage representational structure is itself an efficient response to combinatorial complexity. In natural environments, behavior depends on conjunctions of perceptual features interpreted under changing goals and contexts, so storing a separate representation for every possible stimulus–goal combination would scale poorly and limit generalization to novel situations [13, 78–80]. Representing reusable perceptual primitives instead allows the same sensory components to be recombined flexibly across tasks, with more abstract and behaviorally relevant variables constructed compositionally from those parts [79, 81]. Sequential transformations across multiple stages from perceptual descriptions to variables such as value may therefore reflect not just a descriptive feature of cognition, but an efficient architecture for flexible behavior under resource constraints. Consistent with this view, our results show that behavior is best explained by models in which representations are constructed sequentially across multiple stages, and within this architecture resource limitations further shape the representations formed at each stage. Efficiency principles may thus operate at two levels: the structure of the representational system, and the representations it produces.

A further contribution of this work lies in the experimental leverage provided by the paradigm itself. By holding perceptual input statistics fixed while manipulating the stimulus–value mapping, we fully specified the critical priors and mappings through the task, leaving stage-specific noise as the only free parameter(s). This yields qualitative predictions about the direction of biases across orientations and conditions that differ sharply across representational architectures, so model dissociation does not rely on fine parameter tuning but on the presence or absence of predicted qualitative signatures. The same logic could be applied to other stimulus domains in which perceptual quantities are mapped onto abstract decision variables. Numerical magnitude and numerosity, for example, exhibit systematic biases consistent with efficient coding and Bayesian decoding [82–85], and in many economic contexts such quantities are mapped onto values such as monetary rewards. Similar paradigms could therefore help dissociate how distortions originating in magnitude representations, stimulus–value mappings, and value priors jointly shape value-based decisions. A related direction concerns metacognitive variables such as confidence and error monitoring, increasingly understood as abstract quantities constructed from internal estimates of uncertainty and error [86, 87]. Multi-stage representation frameworks like the one studied here may help determine whether metacognitive capacity itself exhibits analogous structure-dependent distortions arising from the representations and transformations that generate it. More broadly, the present study provides a tractable methodological approach for orthogonally manipulating priors across linked representational stages: scalar perceptual inputs are held fixed while their mapping onto scalar value outputs is varied, thereby changing value priors independently of perceptual priors. This logic should extend to other cognitive domains in which behavior depends on abstract, constructed representations rather than raw sensory input alone, and may help establish when common representational principles generalize beyond perception and valuation.

Several limitations of the present study define useful directions for future work. First, although Phases 1 and 2 were designed to ensure thorough learning of the orientation–value mapping before the no-feedback test phase, the decisions participants made during Phase 2 were themselves subject to the same perceptual and valuation processes studied here. Feedback allowed systematic correction of these biases during learning, yet the extent to which the internally represented mapping is fully veridical after training remains unknown. Importantly, this is not a confound but a natural consequence of the framework: if efficient coding and Bayesian decoding govern value-based behavior, the same principles likely shape the learning process through which stimulus–value associations are acquired, an interesting direction for future work. Second, the present study employed a between-subjects design, leaving open whether participants could sequentially learn all three mappings and whether the same effects would emerge within subjects. A within-subject design would provide stronger causal evidence and allow direct comparison of individual bias profiles across mapping conditions. Third, while the present findings establish multi-stage efficient coding and demonstrate its consequences for valuation of orientation stimuli, the extent to which it generalizes across perceptual domains remains open. We deliberately chose orientation for its well-characterized long-term environmental statistics and strong nonlinear biases. Identifying an equivalently diagnostic mapping in other domains would first require working out, via the value prior formula 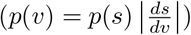 and model simulations, what stimulus–value relationship would generate the necessary qualitative signatures, a question future studies should address.

In sum, our results show that systematic behavioral distortions can arise from multi-stage constructed representations operating under noise and resource constraints. By experimentally dissociating the statistical structure shaping perceptual and value representations, we show that efficient coding and Bayesian decoding can jointly govern both stages, with transformations between them further shaping the geometry of internal noise and the resulting biases. An important next step is to examine how such multi-stage representations are implemented in neural systems. Our results imply multi-stage organization at the level of representations, but whether this corresponds to modular neural architectures or arises from distributed network dynamics remains open. Combining paradigms like the one developed here with neural measurements could help determine how perceptual and value representations are encoded and transformed across stages, and how resource constraints shape their activity. More broadly, the experimental strategy introduced here offers a framework for probing how representations at multiple task-relevant stages are formed and transformed in the brain, helping to link perceptual coding, valuation, and higher cognition through shared representational principles.

## Methods

### Participants

A total of 123 healthy volunteers were recruited through the UAST participant recruitment platform at the University of Zurich between 3–5 December 2025. Eligible participants were at least 18 years old, fluent in English, reported normal or corrected-to-normal vision, and had no prior participation in studies involving deception. Absence of astigmatism was assessed by self-report during recruitment due to the orientation-based nature of the stimuli. Following our preregistered criteria (OSF preregistration: https://osf.io/fcnam/overview), participants were excluded if the ordinary least-squares slope of submitted value on true value fell outside the interval 0.7–1.3 in either Phase 2 or Phase 3. On this basis, four participants were excluded, yielding a final analyzed sample of 119 participants.

Participants were randomly assigned to one of three between-subject orientation– value mapping conditions: *Uniform* (*n* = 39), *Aligned* (*n* = 38), or *Misaligned* (*n* = 42). Mean age was comparable across conditions (Uniform: 23.28 ± 5.20 years; Aligned: 22. ± 97 3.58 years; Misaligned: 23.60 ± 3.15 years; mean ± s.d.). Gender distribution was balanced (Uniform: 22 male, 17 female; Aligned: 21 male, 17 female; Misaligned: 22 male, 20 female). The preregistered target sample size was at least 35 analyzed participants per condition after exclusions. All sessions were conducted in the UZH BLU behavioral laboratory. Participants provided written informed consent prior to participation. The study conformed to the Declaration of Helsinki and was approved by the Human Subjects Committee at the Faculty of Economics, Business Administration, and Information Technology at the University of Zurich. Participants received a base payment of 10 CHF plus choice-dependent compensation determined by a Becker–DeGroot–Marschak procedure in Phase 3 [88], resulting in total earnings of approximately 30–35 CHF. Each session lasted up to 2 hours, and participants typically finished in between 1 hour 15 minutes and 2 hours, depending on how long they took to complete the self-paced Phase 1 and read the instructions.

### Experimental design

The study employed a between-subject design with one primary experimental factor: orientation–value mapping (Uniform, Aligned, Misaligned). Data were collected across four laboratory sessions on three consecutive days. In the first session, all three mapping scripts were loaded simultaneously on separate workstations and participants were randomly allocated to an available workstation upon arrival, determining their mapping condition. In the three subsequent sessions, each session was preconfigured with a single mapping condition, with the mapping assignment randomized across sessions. Participants were unaware of their assigned condition and unaware that other mapping conditions existed.

Across all conditions, stimulus orientations were sampled uniformly throughout the experiment, while the orientation–value mapping differed between conditions according to preregistered functions (Supplementary Table 1). Each mapping was defined over the full orientation domain (0°–180°) and spanned a theoretical value range of 2–42 CHF at these endpoints. Because only orientations between 7.5° and 172.5° were actually presented, however, the realized value range fell short of these extremes and differed across mappings (Supplementary Table 1). All values were rounded to the nearest 0.5 CHF.

Participants completed three sequential phases: observational learning, supervised mapping training, and valuation testing. In each phase, orientations were sampled uniformly within the phase-specific range; Phase 3 used a narrower range centered on the orientations most diagnostic for the predicted effects. Thus, perceptual input statistics were always uniform across mapping conditions, and only the stimulus–value mapping was manipulated across participants, inducing different short-term priors in value space while leaving perceptual priors unchanged. The experiment was implemented in PsychoPy (version 2024.2.1).

### Stimuli and apparatus

Stimuli were Gabor patches displayed on a 24.5-inch monitor (1920 × 1080 pixels, 60 Hz refresh rate) against a grey background. Gabors were circular (diameter: 5° of visual angle; spatial frequency: 3 cycles per degree; random phase on each trial). The intended viewing distance was 80 cm with monitor width of approximately 50 cm; no head stabilization was used and viewing distance was not rigidly enforced, so stimulus size in degrees of visual angle is approximate. Contrast was set to 100% during Phases 1 and 2 to allow accurate encoding of the orientation–value mapping, and reduced to 2% in Phase 3 to introduce additional perceptual uncertainty. Stimulus timing likewise varied by phase, while being identical across the three mapping conditions: viewing was self-paced in Phase 1, whereas Phases 2 and 3 used fixed response windows of 10 s and 4.5 s, respectively, with the Gabor remaining on screen throughout the window (full timing under Procedure). In Phases 2 and 3, each trial began with a 0.5 s fixation cross before stimulus onset; Phase 1 was self-paced and included no fixation cross.

### Procedure

Participants completed three sequential phases: (1) observational learning, (2) supervised mapping training with feedback, and (3) a valuation task without feedback.

#### Phase 1: Observational learning

Participants were introduced to the full orientation–value mapping associated with their assigned condition. High-contrast (100%) Gabor stimuli were presented at 23 discrete orientations ranging from 7.5° to 172.5° in 7.5° increments, each displayed together with its corresponding numerical value. Each orientation–value pairing was displayed for up to 10 s; participants could advance to the next pairing or return to the previous one at any time by pressing ENTER or BACKSPACE respectively. No responses were required in this phase.

#### Phase 2: Learning with feedback

Participants actively responded with the value associated with each of the 23 orientations (100% contrast). On each trial, a fixation cross was presented for 0.5 s, followed by the Gabor stimulus. Participants entered their value using the keyboard (numeric keys with decimal input) and confirmed their response by pressing ENTER. The response window was 10 s. Correct value feedback was then displayed for up to 3 s (or until keypress). Each orientation was presented 10 times in fully randomized order (230 trials total), so all orientation– value pairs received identical exposure during active learning. Orientation sampling was uniform. Thus, any mapping-dependent effects observed in the subsequent no-feedback valuation phase cannot be attributed to differential exposure to particular orientations or values during learning. Phase 2 provided feedback-driven practice on the stimulus–value mapping after participants had already explored the full mapping in Phase 1.

#### Phase 3: Valuation task without feedback

The testing phase consisted of 204 trials. Stimuli were presented at reduced contrast (2%) to increase perceptual uncertainty. Orientations were restricted to 17 values between 30° and 150° in 7.5° increments, each repeated 12 times in fully randomized order. Each trial began with a 0.5 s fixation cross, followed by the Gabor stimulus. Participants had 4.5 s to submit a monetary bid that ideally should correspond to the value they believed the orientation carried according to the learned mapping (in order to maximize potential earnings). The Gabor stimulus remained visible throughout the response window. Responses were entered via keyboard and confirmed with ENTER. No feedback was provided. Orientation sampling remained uniform throughout this phase.

#### Incentive structure

Phase 3 used a Becker–DeGroot–Marschak (BDM) auction mechanism [88] to elicit participants’ subjective value for each presented orientation. The BDM is incentive-compatible: under this mechanism, a participant maximizes expected earnings by bidding exactly their subjective valuation of the item, so bids provide an incentive-compatible readout of internal value estimates. On each Phase 3 trial, participants were endowed with a trial budget of 42 CHF and submitted a bid *b* for the presented stimulus. A random price *p* was then drawn uniformly between 2 and 42 CHF. If the bid was at least as high as the random price (*b* ≥ *p*), the participant purchased the stimulus at price *p* and earned the remaining budget plus the stimulus’s learned monetary value *v*, yielding trial payoff 42 *− p* + *v*. If the bid was lower than the random price (*b < p*), no purchase occurred and the participant retained the full trial budget, yielding trial payoff 42. Total Phase 3 earnings were summed across trials and converted to real payment at a fixed rate of 1/400, in addition to the 10 CHF show-up fee.

### Exclusion criteria and data preprocessing

Exclusion criteria were defined a priori in the preregistration (OSF: https://osf.io/fcnam/overview) and applied without modification.

#### Participant-level exclusions

Participants were excluded if they failed to demonstrate adequate learning of the orientation–value mapping in either Phase 2 (supervised mapping) or Phase 3 (valuation task). Specifically, for each participant and phase, we estimated the ordinary least-squares slope of submitted values (or bids) regressed on the corresponding true values. Participants were excluded if the estimated slope fell outside the preregistered interval [0.7, 1.3] in either phase. Participants were additionally excluded if they provided fewer than 80 valid responses in either Phase 2 (out of 230 total trials in phase 2) or Phase 3 (out of 204 total trials in phase 3).

Applying these criteria resulted in the exclusion of four participants due to slope values outside the preregistered interval. No participants were excluded for insufficient trial counts.

#### Trial-level preprocessing

After participant-level exclusions, trial-level data were cleaned by removing extreme response outliers using the following procedure. For each participant and task phase separately, we computed the distribution of signed errors (submitted value minus true value). Trials whose error deviated by more than three standard deviations from the participant-specific mean error were removed. This procedure was applied separately within Phase 2 and Phase 3. Furthermore, trials on which no response was submitted within the allotted response window were also excluded as missing data (prior to outlier trimming). All subsequent analyses were conducted on the cleaned dataset.

#### Statistical analysis

We tested the model predictions using three complementary analyses. First, we compared the qualitative structure of the observed bias functions with the diagnostic predictions of the competing representational models. Second, we conducted preregistered model-free mixed-effects analyses testing the predicted bias signatures around the cardinal orientation. Third, we performed formal computational model comparison by fitting the competing architectures to trial-wise bids and comparing their relative support. All confirmatory hypotheses, analysis windows, and model specifications were defined a priori in the preregistration (OSF: https://osf.io/fcnam/overview) and implemented without modification. Analyses were restricted to Phase 3 (valuation task) trials. The dependent variable was signed valuation bias, defined as submitted bid minus true value (bias_test_). Following the preregistered confirmatory plan, analyses were restricted to a central orientation window of 65°–115°, centered at 90°. Orientation was mean-centered at 90° (*θ*_*c*_ = *θ* − 90) for all regression models. Although the preregistration specified one-sided tests for directional hypotheses, we report two-sided *p*-values throughout; given the magnitude of the observed effects, inference is identical under either convention.

#### Primary preregistered interaction test (H1)

To test the central prediction that changing the stimulus–value mapping selectively reshapes the bias–orientation relationship, we fit the preregistered linear mixed-effects model:

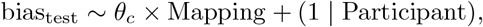

where `Mapping` was a three-level factor (Uniform, Aligned, Misaligned) with Uniform as the reference level. A significant *θ*_*c*_ *×* Mapping interaction confirmed H1.

#### Amplification under Aligned (H3)

Confirmation of H3 required a significantly positive interaction coefficient for the Aligned condition relative to Uniform 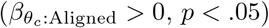, extracted from the primary interaction model.

#### Persistence under Uniform (H2)

To test whether systematic bias persists in the Uniform mapping, reflecting a perceptual contribution independent of short-term value structure, we fit a separate within-condition model restricted to Uniform trials:

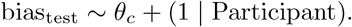

A significantly positive slope of *θ*_*c*_ within Uniform confirmed H2.

#### Slope reversal under Misaligned (H4)

To test the preregistered prediction that bias reverses sign when perceptual and value priors oppose one another, we fit the same within-condition model restricted to Misaligned trials only. A significantly negative slope of *θ*_*c*_ within Misaligned confirmed H4.

#### Aligned–Misaligned contrast

To quantify the predicted amplification versus attenuation of bias magnitude when perceptual and value priors align versus compete, a planned linear contrast was computed from the primary interaction model comparing bias slopes between Aligned and Misaligned mappings, using the delta method to obtain standard errors.

All models were estimated using maximum likelihood (`REML = FALSE`) in `lme4`, with `lmerTest` used to obtain Satterthwaite-approximated degrees of freedom and two-sided *p*-values for fixed effects. Random effects included participant-specific intercepts. All analyses were conducted in R via `rpy2` (Python 3.11.6).

### Response variability

Trial-to-trial response variability was quantified as the within-participant standard deviation of submitted bids at each tested orientation in Phase 3, averaged across participants within each mapping condition. Predicted variability was computed analogously from the model-implied bid distribution under the best-fitting parameters.

### Computational modelling

#### Overview

We formalized three competing representational architectures to test whether efficient coding and Bayesian decoding operate only in perceptual space, only in value space, or across both stages. All models were implemented in Python 3.11 using NumPy 1.25.2 and SciPy 1.11.3. In all models, efficient coding was implemented as a cumulative distribution function (CDF) transformation based on the prior of the represented variable (a standard redundancy-reducing efficient code) [27–29], followed by Bayesian decoding under an explicit loss function. The three architectures differed based on which representational stage(s) were subject to this process.

#### Orientation priors

Orientation was represented on a doubled-angle circular domain, mapping stimulus orientations *θ* ∈ [0, 180) onto *ϕ* ∈ [0, 2*π*) via *ϕ* = 2*θ · π/*180. This reparameterization maps the half-circle of orientations onto a full circle, allowing the use of standard circular statistics and von-Mises distributions which are defined over [0, 2*π*) while preserving the periodicity and symmetry properties of orientation space. All reported orientations are in the original [0, 180) domain; the doubled-angle is only a reparameterization in the code used for model computations.

#### Perceptual encoding and decoding

For a presented orientation *ϕ*_0_, efficient perceptual encoding warped the stimulus domain according to the CDF of the orientation prior *p*_ori_(*ϕ*):

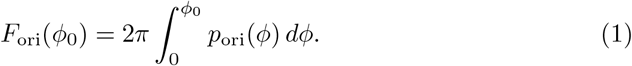

Noise in perceptual representational space was modeled as a von Mises distribution,

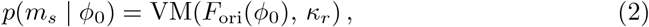

where *κ*_*r*_ is the perceptual precision parameter. Bayesian decoding recovered an internal orientation estimate via

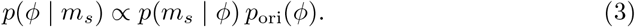

The decoded estimate 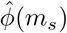 minimized expected circular loss

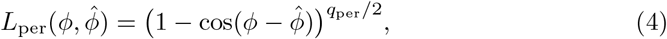

with *q*_per_ = 8 fixed across all fits [55]. For each model variant, the orientation prior used for encoding and decoding was identical within that variant: either the long-term prior *p*_long_ or the short-term uniform prior *p*_short_ for both stages.

The experimenter-observable distribution 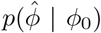 was obtained by propagating *p*(*m*_*s*_ | *ϕ*_0_) through the decoding map 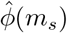 using a mass-conserving numerical pushforward, in which probability mass in each grid bin was redistributed onto the decoded orientation grid via histogram accumulation. This distribution was then pushed through the condition-specific stimulus–value mapping *G* to obtain the perceptually induced distribution over values, 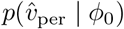.

#### Value priors, encoding, and decoding

The contextual prior over values, *p*_*V*_ (*v*), was derived by pushing the short-term uniform orientation prior through the condition-specific mapping *G*, yielding the distribution of values implied by uniform orientation sampling in each condition. All value computations were performed on a dense grid of 101 points over [*v*_min_, *v*_max_] = [2, 42] CHF. Efficient value encoding warped the value domain according to the CDF of *p*_*V*_ (*v*):

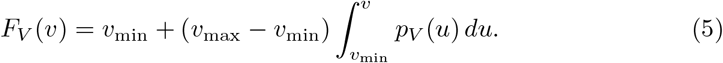

Noise in value representational space was modeled as a Gaussian truncated to the admissible value range:

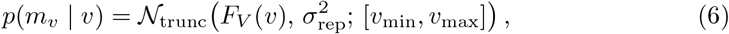

where *σ*_rep_ is the value-noise parameter. Bayesian decoding in value space used

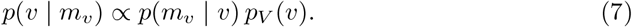

The internal value estimate 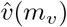minimized posterior expected *q*_val_-norm loss. In all reported fits *q*_val_ = 2, such that decoding corresponded to the posterior mean:

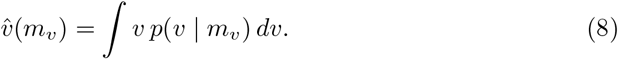

#### Model architectures

Under the *efficient-perception* model, only perceptual representations were subject to efficient coding and Bayesian decoding. The decoded percept 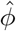 was transformed deterministically into value by the mapping *G*, so that all structure-dependent distortions arose in perceptual space and due to the mapping dependent reshaping of earlier perceptual noise, but not in value space itself:

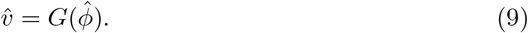

Under the *efficient-valuation* model, perception was veridical at the level of the mapping input as the presented orientation was mapped directly into value space, and efficient coding with Bayesian decoding operated only on the resulting value representation:

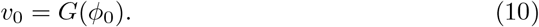

All structure-dependent distortions therefore arose in value space.

Under the *multi-stage* model, perceptual and value representations were both subject to efficient coding and Bayesian decoding. The perceptual model first generated the full distribution of perceptually induced values 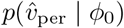. This distribution was then integrated through the value-stage encoder to yield an encoded value distribution:

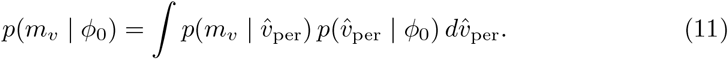

Bayesian decoding in value space then produced the final experimenter-observable distribution 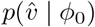. The multi-stage model therefore allowed independent contributions of perceptual and value priors, with perceptual uncertainty propagating forward into value space before additional value-stage inference.

Each of the three architectures was instantiated under two perceptual-prior assumptions: long-term environmental prior or short-term uniform prior for both encoding and decoding. This yields six candidate models for confirmatory model comparison.

#### Descriptive categorical-stabilization extension

To account for the localized collapse in empirical response variability near the 90° cardinal, we fit an additional descriptive extension of the multi-stage model. This extension left perceptual and value encoding and decoding unchanged but applied a hard category gate to the final value-estimate distribution 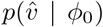. Stimuli were classified into three categories relative to the cardinal: below, at, or above 90°. The near-cardinal category was defined as orientations within one model grid step of 90°. In value space, the middle category corresponded to the interval [*v*_mid_ *−δ, v*_mid_ +*δ*] with *v*_mid_ = 22 CHF and *δ* = 0.25 CHF; lower and upper categories corresponded to values outside this interval. Probability mass outside the implied category was set to zero and the remaining distribution was re-normalized. This extension was descriptive and introduced no additional free parameters. Moreover, the critical preregistered analysis results did not change with this model extension.

### Model fitting and comparison

#### Fit targets and parameter grids

All models were fit separately to each participant’s Phase 3 trial-wise bid data by maximum likelihood. The perceptual noise parameter *κ*_*r*_ was evaluated on a 50-point grid spanning [2, 101], and the value noise parameter *σ*_rep_ was evaluated on a 50-point grid spanning [0.01, 4]. The perceptual and value loss exponents were fixed across all fits at *q*_per_ = 8 and *q*_val_ = 2, respectively. Models included a fixed lapse rate *λ*; primary comparisons used *λ* = 0.01, with robustness checks at *λ* ∈ {0.02, 0.03} (Supplementary Table 2).

#### Trial-wise likelihood

Model fitting was based on the predicted distribution of bids in value space, 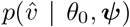, where *θ*_0_ denotes the presented orientation and ***ψ*** denotes the free parameter set of the model. Because responses were recorded on a 0.5-CHF grid, each observed bid was first rounded to the nearest 0.5-CHF grid center. The probability assigned to an observed bid *b*_obs_ was then computed as the model-predicted probability mass in the corresponding response bin of half-width 0.25 CHF:

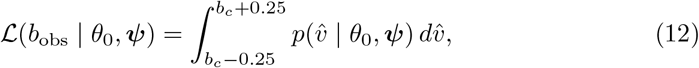

where *b*_*c*_ is the rounded response center. This bin probability was obtained by numerically integrating the model-predicted density via linear interpolation of the cumulative distribution function evaluated on the model grid.

To accommodate occasional random responses, this likelihood was mixed with a lapse component assigning uniform probability mass across the bid domain:

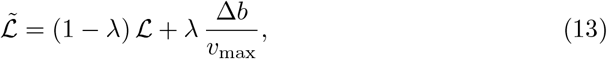

where *λ* is the lapse rate (fixed on 0.01 by default), Δ*b* = 0.5 CHF is the response-bin width and the lapse distribution is uniform over [0, 42] CHF, reflecting that a lapsed response could take any value in the full response range, including below the mapping minimum of *v*_min_ = 2 CHF. The negative log-likelihood for a participant was then

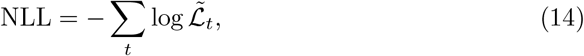

summed over all Phase 3 trials *t*.

#### Optimization

Parameter estimation was performed by exhaustive grid search. For the efficient-perception model, a one-dimensional search was carried out over *κ*_*r*_. For the efficient-valuation model, a one-dimensional search was carried out over *σ*_rep_. For the multi-stage model and the descriptive categorical-stabilization extension, a two-dimensional search was carried out over all (*κ*_*r*_, *σ*_rep_) combinations. For each participant and model, the best-fitting parameter set was defined as the grid point yielding the minimum individual NLL. Fits were parallelized across participants using `joblib`.

#### Parameter recovery

To verify that the grid-search fitting procedure could reliably identify the free parameters of the multi-stage model, we conducted a parameter recovery analysis. Synthetic bid data were generated by sampling from the model-predicted distribution 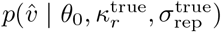 under the same orientation sequences and mapping conditions as the actual Phase 3 data, across a range of true parameter values drawn from the fitting grids. The same grid-search procedure was then applied to recover parameter estimates from the simulated data. Results are reported in Supplementary Figure S1.

#### Model recovery

To verify that the competing models are distinguishable at the group level, at which model comparison was performed, we additionally conducted a group-level model recovery analysis. For each candidate model we generated synthetic Phase 3 datasets (N = 40 simulated participants per mapping, 50 repetitions) using parameters drawn from the fitting grids, refit all six candidates, and selected the best model by group-level AIC summed across simulated participants, exactly as for the empirical data. The generating architecture was recovered in essentially all repetitions (Supplementary Fig. 10).

#### Model comparison

To compare representational architectures at the population level, individual-participant NLLs were summed within each model to obtain a total negative log-likelihood. Model complexity was penalized using the Akaike Information Criterion,

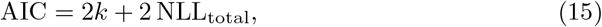

and the Bayesian Information Criterion,

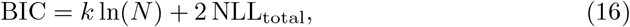

where *k* denotes the number of free parameters for a given participant and model, and *N* denotes that participant’s Phase 3 trial count; individual AIC and BIC values were then summed across participants to obtain population-level scores. Lower AIC and BIC values indicate better model support after penalizing model complexity. Full model comparison results across all architectures, perceptual prior assumptions (long-term and short-term), and lapse rates are reported in Supplementary Table 2.

## Acknowledgments

We thank C. Schnyder and E. Boldin for assistance with recruitment of subjects. This research was supported by the University Research Priority Program (URPP) Adaptive Brain Circuits in Development and Learning (AdaBD) at the University of Zurich.

## Data and code availability

Data and analysis code will be made publicly available upon publication.

## A Supplementary Tables

**Table 1. Supplementary Table 1.** Orientation–value mappings for the three experimental conditions. Discrete orientation–value pairs used in the Uniform, Aligned, and Misaligned conditions. All values are in CHF, rounded to the nearest 0.5 CHF. Orientations are in degrees. The theoretical value range spans 2–42 CHF, corresponding to idealized orientations of 0° and 180°; presented orientations ranged from 7.5° to 172.5° in 7.5° steps. The Uniform mapping assigned values via linear interpolation between 2 and 42 CHF across the full orientation range. The Misaligned mapping used the scaled CDF transformation of the same prior, concentrating values near those associated with oblique orientations (around 45° and 135°), inducing a value prior misaligned with the long-term perceptual prior. The Aligned mapping used an inverted mapping of the scaled-CDF transformation of the long-term orientation prior *p*_long_(*ϕ*) ∝ 2 −|sin *ϕ*|, concentrating values near those associated with cardinal orientations (around 90°), thereby inducing a short-term value prior aligned with the long-term perceptual prior.

| Orientation (°) | Uniform (CHF) | Aligned (CHF) | Misaligned (CHF) |
| --- | --- | --- | --- |
| 7.5 | 3.5 | 5.5 | 2.5 |
| 15 | 5.5 | 8.0 | 3.0 |
| 22.5 | 7.0 | 10.0 | 4.0 |
| 30 | 8.5 | 11.0 | 5.0 |
| 37.5 | 10.5 | 11.5 | 7.5 |
| 45 | 12.0 | 12.0 | 11.5 |
| 52.5 | 13.5 | 12.5 | 16.0 |
| 60 | 15.5 | 13.0 | 18.5 |
| 67.5 | 17.0 | 14.0 | 20.0 |
| 75 | 18.5 | 16.0 | 21.0 |
| 82.5 | 20.5 | 18.5 | 21.5 |
| 90 | 22.0 | 22.0 | 22.0 |
| 97.5 | 23.5 | 25.5 | 22.5 |
| 105 | 25.5 | 28.0 | 23.0 |
| 112.5 | 27.0 | 30.0 | 24.0 |
| 120 | 28.5 | 31.0 | 25.5 |
| 127.5 | 30.5 | 31.5 | 28.0 |
| 135 | 32.0 | 32.0 | 32.5 |
| 142.5 | 33.5 | 32.5 | 36.5 |
| 150 | 35.5 | 33.0 | 39.0 |
| 157.5 | 37.0 | 34.0 | 40.0 |
| 165 | 38.5 | 36.0 | 41.0 |
| 172.5 | 40.5 | 38.5 | 41.5 |

**Table 2. Supplementary Table 2.** Model comparison results. Total negative log-likelihood (NLL), Akaike Information Criterion (AIC), and Bayesian Information Criterion (BIC) for all candidate models, ranked by AIC. MultiCategorical denotes the descriptive categorical-stabilization extension of the multi-stage model, fit only under the long-term perceptual prior. Long-term: perceptual encoding and decoding governed by the long-term environmental orientation prior. Uniform: perceptual encoding and decoding governed by the short-term uniform prior. Lower AIC and BIC indicate better model support after complexity penalization.

| Rank | Lapse | Architecture | Perceptual prior | NLL | AIC | BIC |
| --- | --- | --- | --- | --- | --- | --- |
| *MultiCategorical* |  |  |  |  |  |  |
| 1 | 0.01 | MultiCategorical | long-term | 61435.09 | 123346.18 | 124125.23 |
| 2 | 0.02 | MultiCategorical | long-term | 61500.93 | 123477.86 | 124256.91 |
| 3 | 0.03 | MultiCategorical | long-term | 61607.01 | 123690.03 | 124469.08 |
| *Multi-stage* |  |  |  |  |  |  |
| 4 | 0.01 | Multi-stage | long-term | 64266.30 | 129008.61 | 129787.66 |
| 5 | 0.02 | Multi-stage | long-term | 64367.42 | 129210.85 | 129989.90 |
| 6 | 0.01 | Multi-stage | uniform | 64408.34 | 129292.69 | 130071.74 |
| 7 | 0.03 | Multi-stage | long-term | 64486.47 | 129448.95 | 130227.99 |
| 8 | 0.02 | Multi-stage | uniform | 64514.23 | 129504.45 | 130283.50 |
| 9 | 0.03 | Multi-stage | uniform | 64635.77 | 129747.55 | 130526.60 |
| *Value-only* |  |  |  |  |  |  |
| 10 | 0.01 | Value-only | long-term<sup>a</sup> | 65165.52 | 130569.03 | 130958.56 |
| 11 | 0.01 | Value-only | uniform<sup>a</sup> | 65165.52 | 130569.03 | 130958.56 |
| 12 | 0.02 | Value-only | long-term<sup>a</sup> | 65233.57 | 130705.14 | 131094.66 |
| 13 | 0.02 | Value-only | uniform<sup>a</sup> | 65233.57 | 130705.14 | 131094.66 |
| 14 | 0.03 | Value-only | long-term<sup>a</sup> | 65329.47 | 130896.94 | 131286.46 |
| 15 | 0.03 | Value-only | uniform<sup>a</sup> | 65329.47 | 130896.94 | 131286.46 |
| *Perceptual-only* |  |  |  |  |  |  |
| 16 | 0.01 | Perceptual-only | uniform | 65382.55 | 131003.10 | 131392.62 |
| 17 | 0.02 | Perceptual-only | uniform | 65481.22 | 131200.45 | 131589.97 |
| 18 | 0.03 | Perceptual-only | uniform | 65593.95 | 131425.91 | 131815.43 |
| 19 | 0.01 | Perceptual-only | long-term | 66726.91 | 133691.81 | 134081.34 |
| 20 | 0.02 | Perceptual-only | long-term | 66796.61 | 133831.21 | 134220.74 |
| 21 | 0.03 | Perceptual-only | long-term | 66884.21 | 134006.42 | 134395.95 |

<sup>a</sup>The Value-only architecture treats perception as veridical, rendering the perceptual prior assumption irrelevant; NLL is therefore identical across both prior configurations.

**Table 3. Supplementary Table 3.** Full mixed-effects model results for preregistered confirmatory analyses. All models were estimated by maximum likelihood (`REML = FALSE`) in `lme4` with Satterthwaite-approximated degrees of freedom from `lmerTest`. The primary interaction model (H1, H3) was fit to all Phase 3 trials in the preregistered orientation window (65°–115°; 9,615 observations, 119 participants), with Uniform as the reference level. Within-condition models for H2 and H4 were fit separately to Uniform (3,134 observations, 39 participants) and Misaligned trials (3,400 observations, 42 participants) respectively. The Aligned–Misaligned planned contrast was computed from the primary interaction model using the delta method. All *p*-values are two-sided.

| Test | Predictor | Estimate | SE | *t* | df | *p* |
| --- | --- | --- | --- | --- | --- | --- |
| *Primary interaction model: bias ~ $\theta_c \times$ Mapping + (1 \| Participant)* |  |  |  |  |  |  |
| H1/H3 | Intercept | −0.2004 | 0.0815 | −2.458 | 119.3 | .015 |
| H1/H3 | $\theta_c$ (Uniform slope) | 0.01442 | 0.00220 | 6.570 | 9497 | < .001 |
| H1/H3 | Mapping: Aligned | −0.0800 | 0.1160 | −0.690 | 119.2 | .491 |
| H1/H3 | Mapping: Misaligned | 0.1472 | 0.1132 | 1.301 | 119.2 | .196 |
| H1 | $\theta_c \times$ Aligned | 0.04294 | 0.00313 | 13.739 | 9498 | < .001 |
| H1 | $\theta_c \times$ Misaligned | −0.04016 | 0.00304 | −13.201 | 9498 | < .001 |
| *Planned contrast (from primary model)* |  |  |  |  |  |  |
| — | Aligned – Misaligned slope | 0.08310 | 0.00306 | 27.123 | 9609 | < .001 |
| *Within-Uniform model (H2): bias ~ $\theta_c$ + (1 \| Participant)* |  |  |  |  |  |  |
| H2 | Intercept | −0.2008 | 0.0710 | −2.827 | 38.75 | .007 |
| H2 | $\theta_c$ | 0.01443 | 0.00232 | 6.229 | 3095 | < .001 |
| *Within-Misaligned model (H4): bias ~ $\theta_c$ + (1 \| Participant)* |  |  |  |  |  |  |
| H4 | Intercept | −0.05294 | 0.08768 | −0.604 | 42.10 | .549 |
| H4 | $\theta_c$ | −0.02573 | 0.00199 | −12.921 | 3359 | < .001 |

**Table 4. Supplementary Table 4.** Best-fitting parameter estimates for all candidate models. Mean and standard deviation (SD) of best-fitting parameters across participants, reported separately by model architecture, perceptual prior assumption, and mapping condition. *κ*_*r*_: perceptual precision (von Mises concentration); *σ*_rep_: value noise. Dashes indicate parameters not estimated under that architecture. All fits used lapse rate *λ* = 0.01; model comparison across lapse rates *λ* ∈ {0.01, 0.02, 0.03} is reported in Supplementary Table 2. Long-term: perceptual encoding governed by long-term environmental orientation prior. Uniform: perceptual encoding governed by short-term uniform prior.

| Architecture | Perceptual prior | Condition | $N$ | $\kappa_r$ | | $\sigma_{\text{rep}}$ | |
| --- | --- | --- | --- | --- | --- | --- | --- |
|  |  |  |  | Mean | SD | Mean | SD |
| Perceptual-only | Long-term | Uniform | 39 | 12.15 | 6.85 | — | — |
|  |  | Aligned | 38 | 19.92 | 14.61 | — | — |
|  |  | Misaligned | 42 | 13.21 | 13.01 | — | — |
| Perceptual-only | Uniform | Uniform | 39 | 9.82 | 4.91 | — | — |
|  |  | Aligned | 38 | 15.61 | 10.77 | — | — |
|  |  | Misaligned | 42 | 10.42 | 9.56 | — | — |
| Value-only | —<sup>a</sup> | Uniform | 39 | — | — | 2.26 | 0.56 |
|  |  | Aligned | 38 | — | — | 1.99 | 0.75 |
|  |  | Misaligned | 42 | — | — | 2.28 | 0.76 |
| Multi-stage | Long-term | Uniform | 39 | 61.73 | 35.17 | 1.90 | 0.56 |
|  |  | Aligned | 38 | 23.69 | 20.68 | 0.32 | 0.44 |
|  |  | Misaligned | 42 | 63.29 | 37.56 | 1.98 | 0.87 |
| Multi-stage | Uniform | Uniform | 39 | 20.65 | 26.60 | 0.70 | 0.82 |
|  |  | Aligned | 38 | 27.04 | 29.58 | 0.65 | 0.81 |
|  |  | Misaligned | 42 | 75.89 | 39.36 | 2.00 | 0.93 |
| MultiCategorical | Long-term | Uniform | 39 | 82.66 | 31.91 | 2.19 | 0.55 |
|  |  | Aligned | 38 | 24.65 | 25.42 | 0.42 | 0.64 |
|  |  | Misaligned | 42 | 67.09 | 38.97 | 2.13 | 0.82 |

<sup>a</sup>The Value-only architecture treats perception as veridical; the perceptual prior assumption is irrelevant.

## Supplementary Figures

### Parameter recovery

We generated synthetic bid data from the model-predicted distribution across the fitting grids (*κ*_*r*_ ∈ [2, 101]; *σ*_rep_ ∈ [0.01, 3.5]), using the same trial structure as Phase 3. The same grid-search procedure was then applied to recover parameters from the simulated data. Recovery performance was quantified as the Pearson correlation between true and recovered parameter values and is shown separately for each model and condition in Supplementary Figures S1–S3. Across models and parameters, recovery correlations ranged from moderate to high.

**Supplementary Figure 1:**
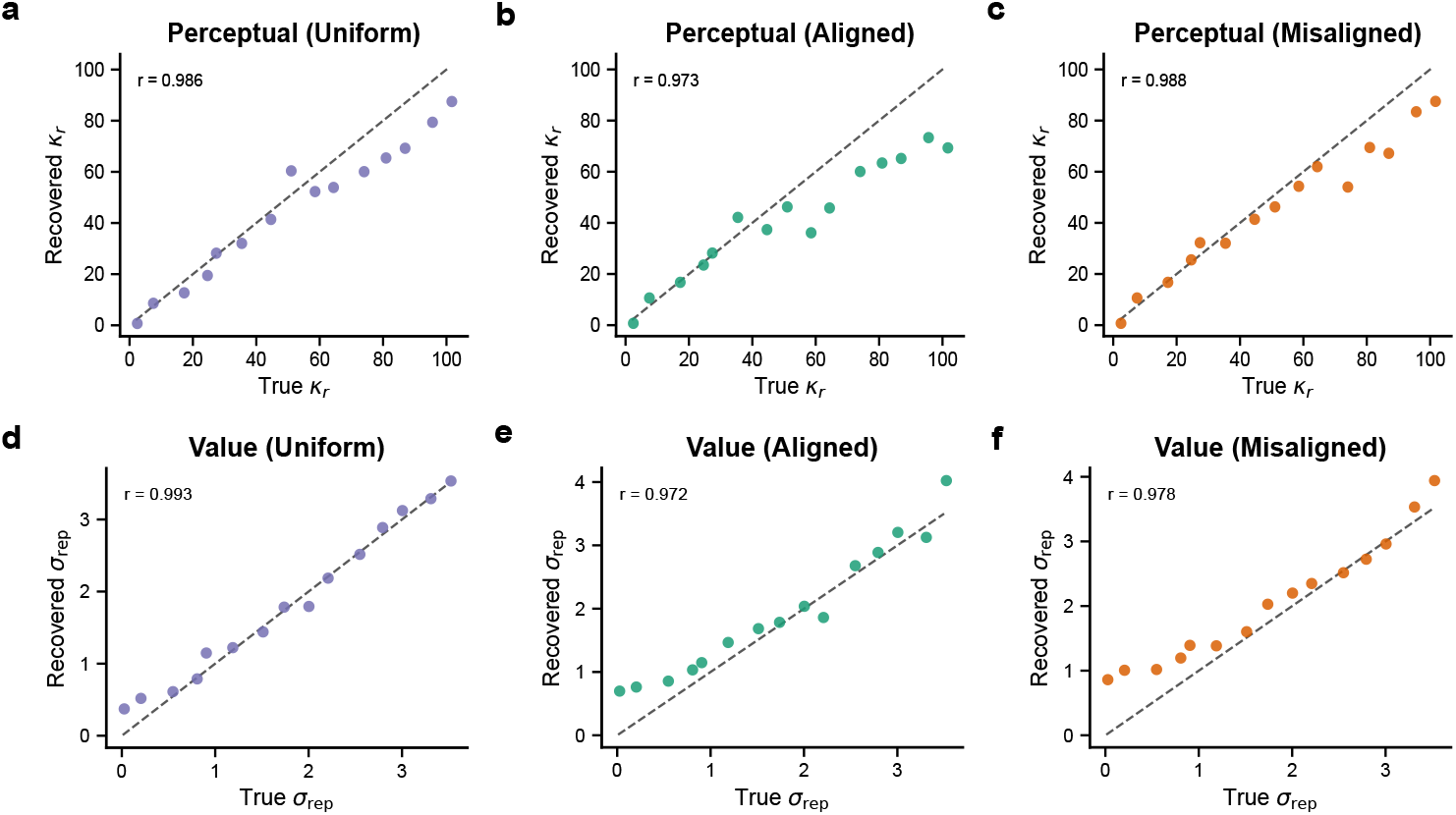
Parameter recovery for single-stage models. Each panel shows recovered versus true parameter values from synthetic data, with the dashed line indicating perfect recovery. (**a–c**) Recovery of the perceptual precision parameter *κ*_*r*_ for the Perceptual-only model across Uniform, Aligned, and Misaligned mapping conditions. Recovery was *r ≥* 0.973 across conditions. (**d–f**) Recovery of the value noise parameter *σ*_rep_ for the Value-only model across conditions. Recovery was *r ≥* 0.972 across conditions.

**Supplementary Figure 2:**
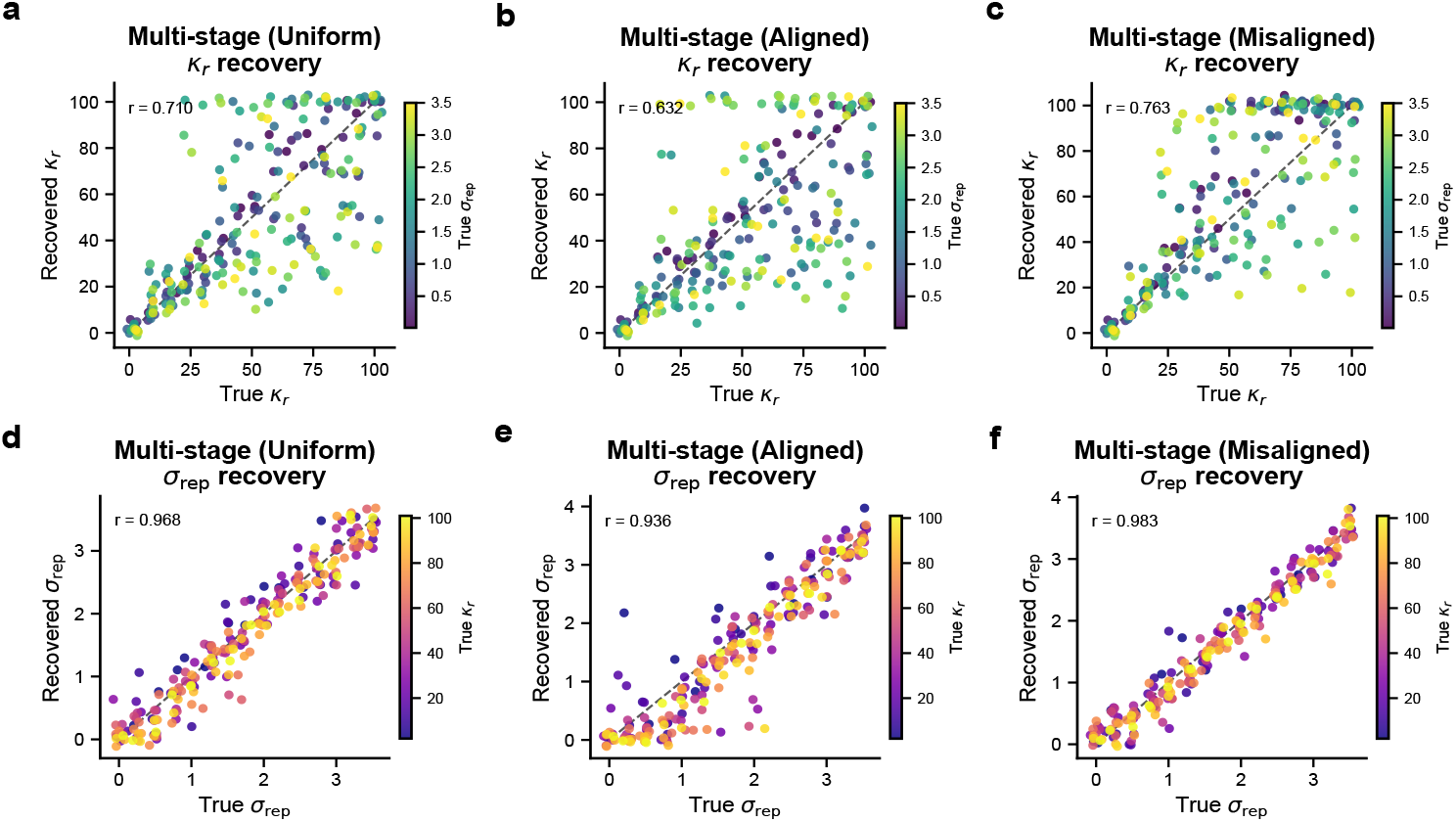
Parameter recovery for multi-stage model. Each panel shows recovered versus true parameter values from synthetic data, with the dashed line indicating perfect recovery. (**a–c**) Recovery of *κ*_*r*_ across Uniform, Aligned, and Misaligned conditions. Recovery was *r* = 0.710 for Uniform, *r* = 0.632 for Aligned, and *r* = 0.763 for Misaligned. (**d–f**) Recovery of *σ*_rep_ across conditions. Dot color indicates true *κ*_*r*_. Recovery was *r* ≥ 0.936 across conditions.

**Supplementary Figure 3:**
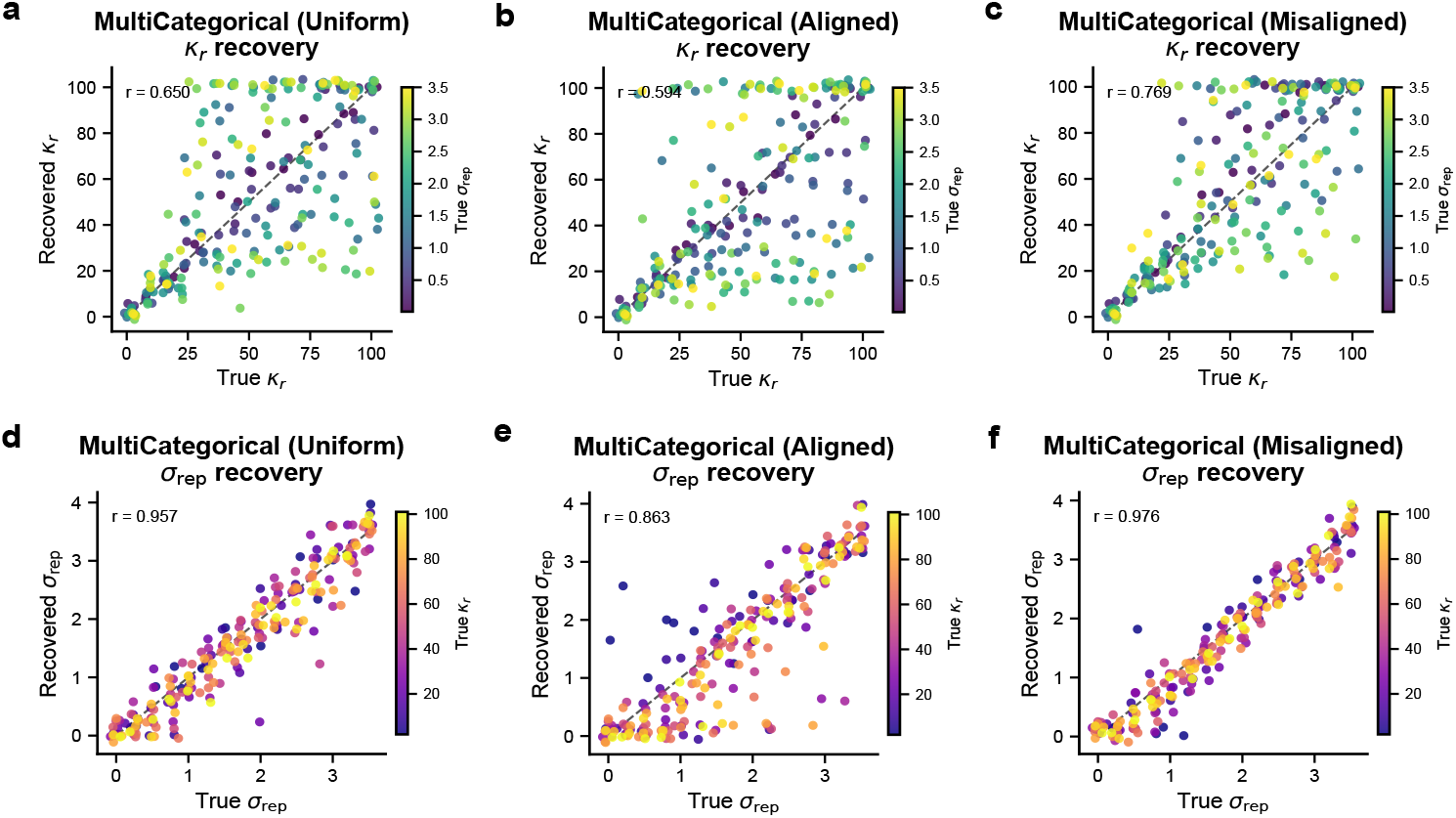
Parameter recovery for Categorical multi-stage model. Each panel shows recovered versus true parameter values from synthetic data, with the dashed line indicating perfect recovery. (**a–c**) Recovery of *κ*_*r*_ across Uniform, Aligned, and Misaligned conditions. Recovery was *r* = 0.650 for Uniform, *r* = 0.594 for Aligned, and *r* = 0.769 for Misaligned. (**d–f**) Recovery of *σ*_rep_ across conditions. Dot color indicates true *κ*_*r*_. Recovery was *r* ≥ 0.863 across conditions.

**Fig. 10.**
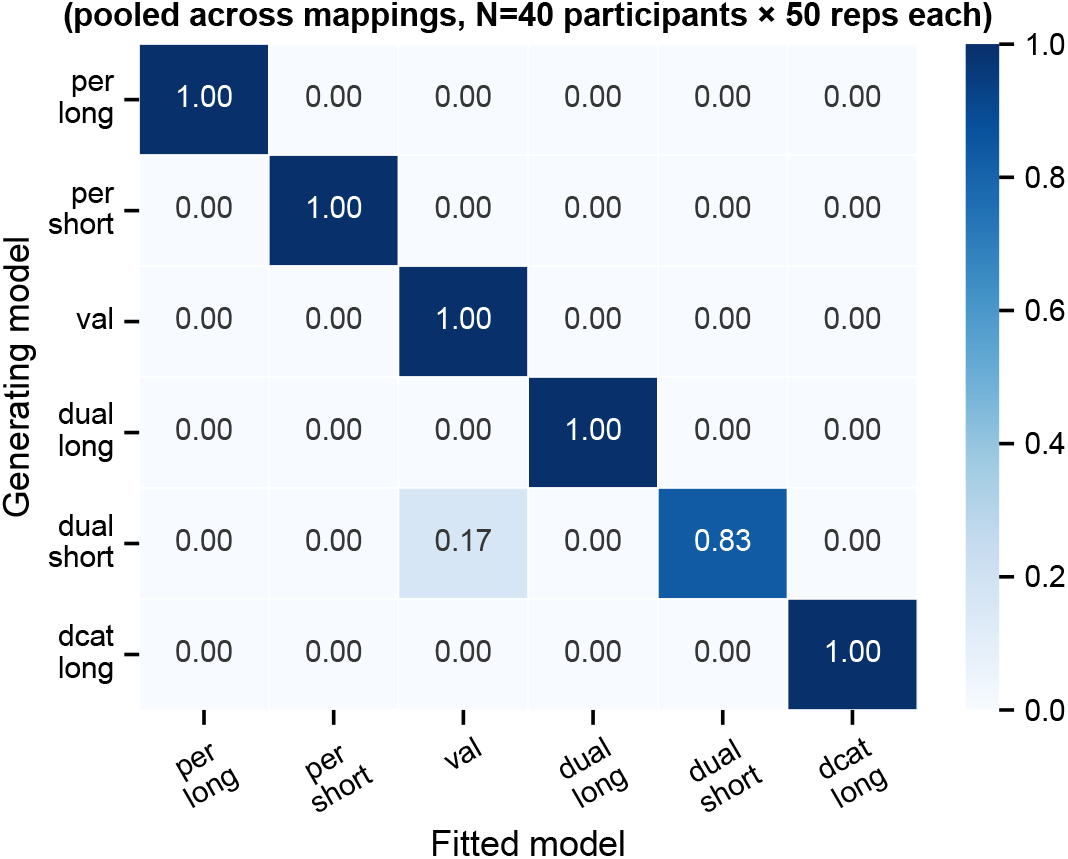
Group-level model recovery. Confusion matrix showing recovery of the generating model architecture at the group level. Synthetic datasets were generated from each of the six candidate models across the three mapping conditions (N = 40 simulated participants per mapping, 50 repetitions), with generating parameters drawn from the fitting grids. For each repetition, all six models were fit to the synthetic data and compared using group-level AIC summed across simulated participants, exactly as for the empirical fits. Rows denote the generating model and columns the model selected as best; cells are row-normalized, so each row sums to 1 and the diagonal gives the proportion of repetitions in which the generating architecture was correctly recovered. Recovery is near-perfect for all models. These results confirm that the three mapping conditions jointly render the competing representational architectures identifiable from Phase 3 behaviour at the group level.

